# Eastern Equine Encephalitis Virus Rapidly Infects and Disseminates in the Brain and Spinal Cord of Infected Cynomolgus Macaques Following Aerosol Challenge

**DOI:** 10.1101/2020.12.21.423836

**Authors:** Janice A Williams, Simon Y Long, Xiankun Zeng, Kathleen Kuehl, April M Babka, Neil M Davis, Jun Liu, John C Trefry, Sharon Daye, Paul R Facemire, Patrick L Iversen, Sina Bavari, Margaret L Pitt, Farooq Nasar

**Affiliations:** Pathology Division, United States Army Medical Research Institute of Infectious Diseases, 1425 Porter Street, Frederick, Maryland, USA; Virology Division, United States Army Medical Research Institute of Infectious Diseases, 1425 Porter Street, Frederick, Maryland, USA; Therapeutics Division, United States Army Medical Research Institute of Infectious Diseases, 1425 Porter Street, Frederick, Maryland, USA; Office of the Commander, United States Army Medical Research Institute of Infectious Diseases, 1425 Porter Street, Frederick, Maryland, USA

**Author notes:** J.A.W. and S.Y.L. contributed equally to this work. Corresponding authors: Farooq Nasar, Mailing address: United States Army Medical Research Institute of Infectious, Diseases, 1425 Porter Street, Frederick, MD 21702., Margaret L. Pitt, Mailing address: United States Army Medical Research Institute of Infectious Diseases, 1425 Porter Street, Frederick, MD 21702.

## Abstract

Eastern equine encephalitis virus (EEEV) is mosquito-borne virus that produces fatal encephalitis in humans. We recently conducted a first of its kind study to investigate EEEV clinical disease course following aerosol challenge in a cynomolgus macaque model utilizing the state of the art telemetry to measure critical physiological parameters. Following challenge, all parameters were rapidly and profoundly altered, and all nonhuman primates (NHPs) met the euthanasia criteria. In this study, we performed the first comprehensive pathology investigation of tissues collected at euthanasia to gain insights into EEEV pathogenesis. Viral RNA and proteins as well as microscopic lesions were absent in the visceral organs. In contrast, viral RNA and proteins were readily detected throughout the brain including autonomic nervous system (ANS) control centers and spinal cord. However, despite presence of viral RNA and proteins, majority of the brain and spinal cord tissues exhibited minimal or no microscopic lesions. The virus tropism was restricted primarily to neurons, and virus particles (~61-68 nm) were present within axons of neurons and throughout the extracellular spaces. However, active virus replication was absent or minimal in majority of the brain and was limited to regions proximal to the olfactory tract. These data suggest that EEEV initially replicates in/near the olfactory bulb following aerosol challenge and is rapidly transported to distal regions of the brain by exploiting the neuronal axonal transport system to facilitate neuron-to-neuron spread. Once within the brain, the virus gains access to the ANS control centers likely leading to disruption and/or dysregulation of critical physiological parameters to produce severe disease. Moreover, the absence of microscopic lesions strongly suggests that the underlying mechanism of EEEV pathogenesis is due to neuronal dysfunction rather than neuronal death. This study is the first comprehensive investigation of EEEV clinical disease course and pathogenesis in a NHP model and will provide significant insights into the evaluation of countermeasure.

**Author Summary:** EEEV is an arbovirus endemic in parts of North America and is able to produce fatal encephalitis in humans and domesticated animals. Despite multiple human outbreaks during the last 80 years, there are still no therapeutic or vaccines to treat or prevent human disease. One critical obstacle in the development of effective countermeasure is the lack of insights into EEEV pathogenesis in a susceptible animal host. We recently conducted a study in cynomolgus macaques to investigate the disease course by measuring clinical parameters relevant to humans. Following infection, these parameters were rapidly and profoundly altered leading to severe disease. In this study, we examined the potential mechanisms that underlie pathogenesis to cause severe disease. The virus was present in many parts of the brain and spinal cord, however, little or no pathological lesions as well as active virus replication were observed. Additionally, neurons were the predominant target of EEEV and virus transport was facilitated by axonal transport system to spread neuron-to-neuron throughout the brain and spinal cord. These data show that EEEV likely hijacks host cell transport system to rapidly spread in the brain and local/global neuronal dysfunction rather than death is the principal cause of severe disease.

## INTRODUCTION

The genus *Alphavirus* in the family *Togaviridae* is comprised of small, spherical, enveloped viruses with genomes consisting of a single stranded, positive-sense RNA, ~11-12 kb in length. Alphaviruses comprise 31 recognized species and the vast majority utilize mosquitoes as vectors for transmission into vertebrate hosts [1–6]. Mosquito-borne alphaviruses can spillover into the human population and cause severe disease. Old World alphaviruses (chikungunya, o’nyong-nyong, Sindbis, and Ross River) can cause disease characterized by rash and debilitating arthralgia, whereas New World viruses [eastern, western, and Venezuelan equine encephalitis virus] can cause fatal encephalitis.

Eastern equine encephalitis virus (EEEV) is an important pathogen of medical and veterinary importance in North America. EEEV is endemic in the eastern United States and Canada, and the Gulf coast of the United States. The main transmission cycle is between passerine birds and *Culiseta melanura* mosquitoes. However, this cycle can spillover into humans and domesticated animals and cause severe disease with human and equid case-fatality rates of 30-90% and >90%, respectively [6, 7]. Human survivors can suffer from debilitating and permanent long-term neurological sequelae with rates of 35-80% [6, 7]. In addition to natural infections, EEEV was developed as a biological weapon during the cold war by the U.S. and the former Union of Soviet Socialist Republics (USSR). Currently, there are no licensed therapeutics and/or vaccines to prevent or treat EEEV infection and the U.S. population remains vulnerable to natural disease outbreaks and/or bioterrorism events.

In order to develop effective vaccine and therapeutic countermeasures, nonhuman primate (NHP) models have been utilized to recapitulate various aspects of human disease, as well as, to gain insight into viral pathogenesis. We recently conducted a study in cynomolgus macaques to explore EEEV disease course utilizing advance telemetry following aerosol challenge. All physiological parameters observed including temperature, respiration, activity, heart rate, blood pressure, electrocardiogram (ECG), and electroencephalography (EEG) were considerably altered post-challenge for the duration of ~24-100 hrs. In addition, all NHPs exhibited profound disruption of the circadian rhythm, sleep, and food/fluid intake. Accordingly, all NHPs met the euthanasia criteria by ~106-140 hours post-infection (hpi). In this study, we performed a detailed investigation of visceral organs, brain, and spinal cord harvested at euthanasia to gain insights into EEEV pathogenesis.

## RESULTS

### EEEV Associated Pathology in the Visceral Organs

Visceral organs including the heart, liver, lung, kidney, and spleen were collected from NHPs at the time of euthanasia and were examined for virus and/or host induced pathology (Supp. Table 1). There were no EEEV associated necrotic and/or inflammatory lesions in the visceral organs of any of NHPs (Figure 1, Supp. Table 1). In addition, *in situ* hybridization (ISH) and immunohistochemistry (IHC) were unable to detect presence of viral RNA or proteins in any organs, respectively (Supp. Figures 1 and 2).

**Figure 1.**
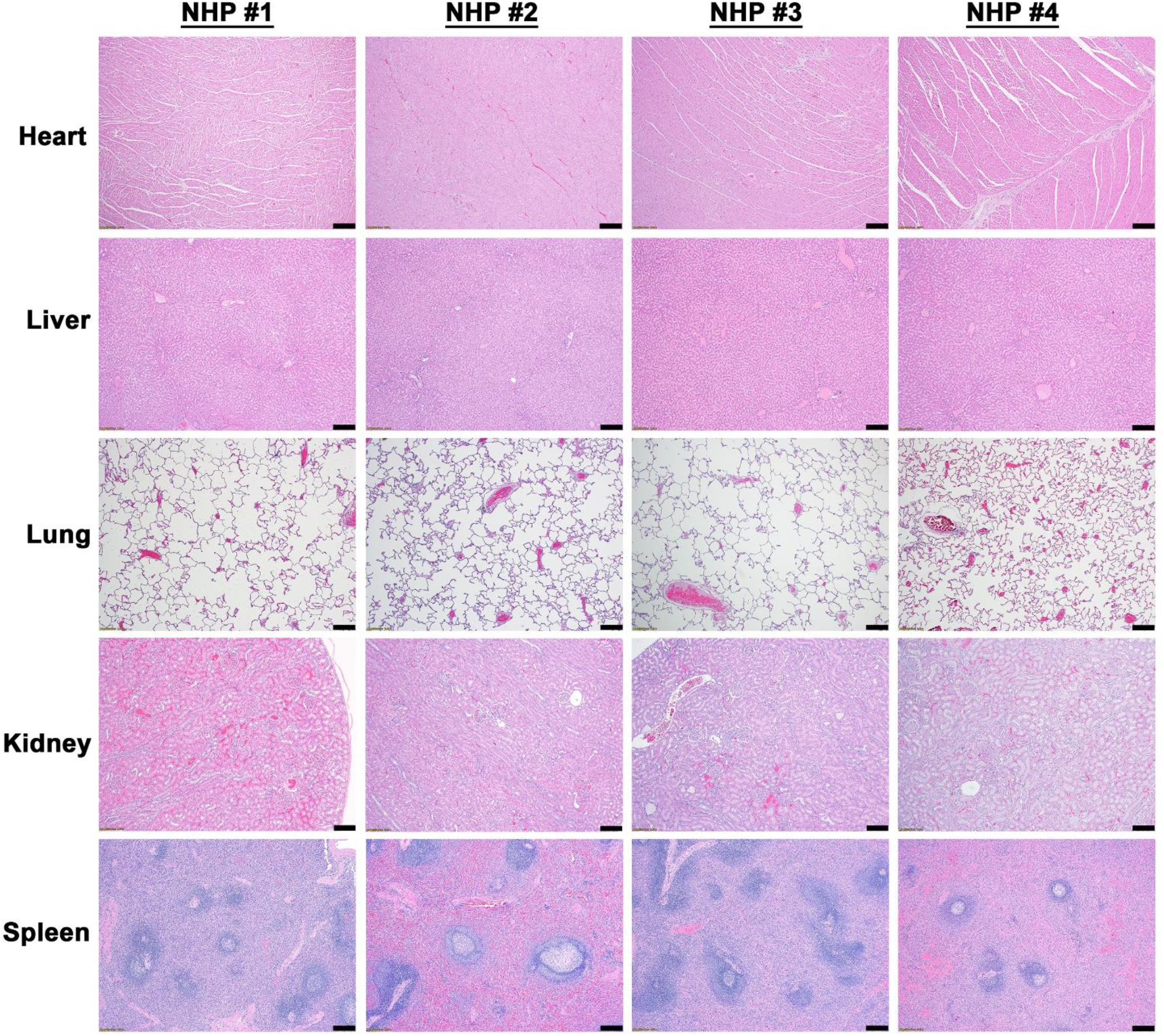
Histopathology in the visceral organs of EEEV infected cynomolgus macaques. The tissues were collected at the time of euthanasia. Hematoxylin and eosin (H&E) staining was performed on the tissues of all four NHPs. Bar = 200 um.

### EEEV Associated Pathology in the Brain and Spinal Cord

The projections from the olfactory bulb connect to the amygdala and hippocampus via the primary olfactory cortex. Our companion manuscript reported the presence of infectious virus in the olfactory bulb of NHPs at the time of euthanasia with titers ranging from 4.1-7.9 log_10_ PFU/mL. Accordingly, the amygdala and hippocampus were investigated for virus and/or host induced pathology. Mild to moderate necrotic and inflammatory lesions were observed throughout both regions of the brain in all NHPs (Figure 2). The necrotic lesions were characterized by neuronal degeneration, satellitosis, and necrosis, as well as vacuolation of the neutrophil (Figure 2). The inflammatory lesions comprised predominantly of neutrophilic infiltrates in all NHP sections except the hippocampus of NHP #2. Furthermore, substantial viral RNA and proteins were readily detected in the amygdala and hippocampus of all NHPs (Figure 2).

**Figure 2.**
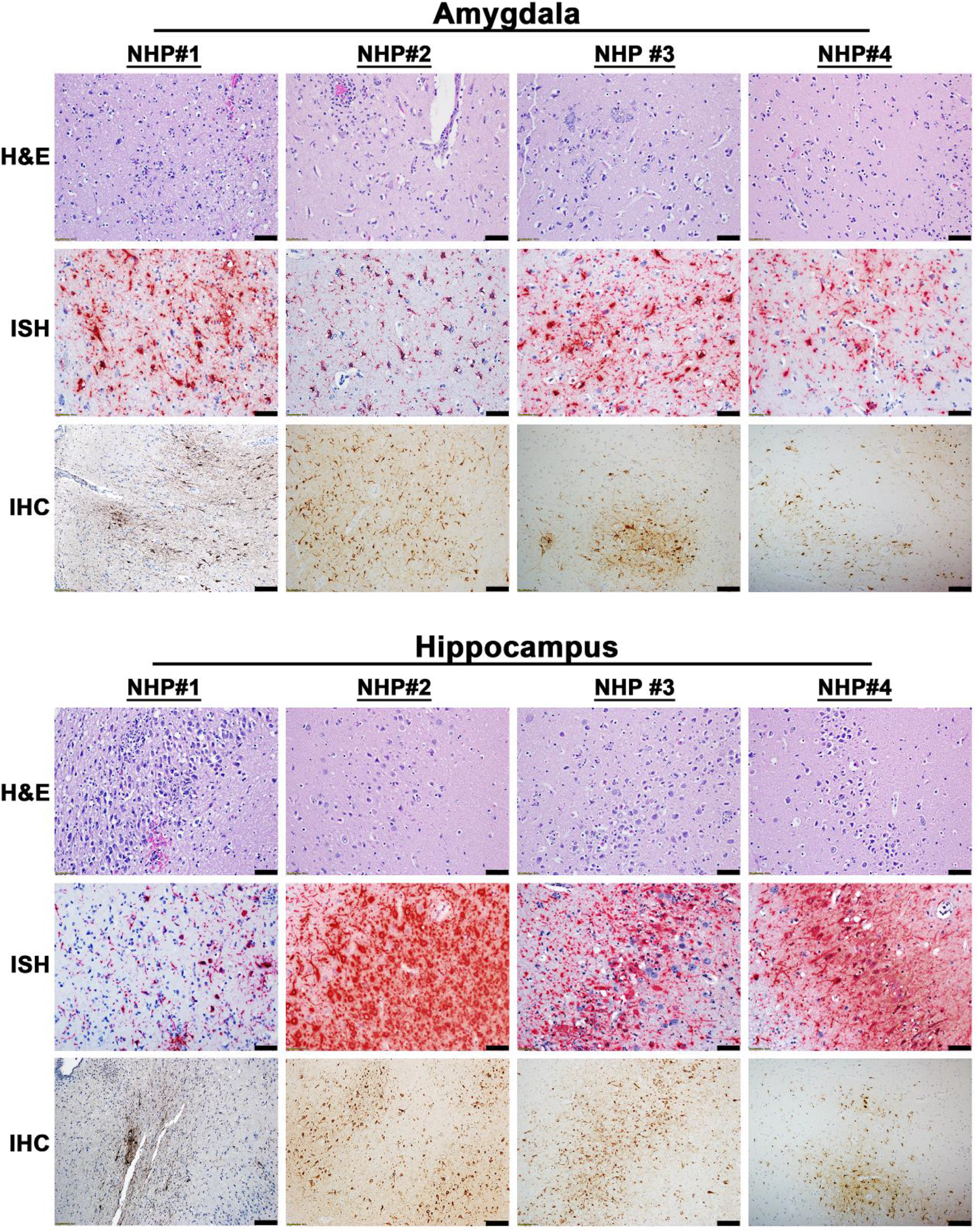
Pathology in the amygdala and the hippocampus of EEEV infected cynomolgus macaques. The tissues were collected at the time of euthanasia. Hematoxylin and eosin (H&E) staining was performed to visualize histopathology. The presence of EEEV RNA and proteins was visualized via *in situ* hybridization (ISH) and immunohistochemistry (IHC), respectively. H&E, ISH, and IHC were performed on the tissues of all four NHPs. Bar = 100 um (H&E and IHC). Bar = 50 um (ISH).

The projection of the amygdala and hippocampus connect to other parts of the midbrain, which in turn are connected to both the forebrain and the hindbrain. We next examined various structures in these regions including the hypothalamus, thalamus, corpus striatum, mesencephalon, medulla oblongata, frontal cortex, and cerebellum for virus and/or host induced pathology (Supp. Table 1). In contrast to the pathology observed in the amygdala and hippocampus, the majority of these tissue sections displayed minimal or no microscopic lesions (Figures 3 and 4). Few focal lesions were observed in some regions and were restricted primarily to the corpus striatum, thalamus, mesencephalon, and medulla oblongata (Figure 4). The focal lesions comprised of minimal to mild neuronal degeneration, necrosis, neuropil vacuolation, gliosis, and neuronal satellitosis (Figure 4). The latter was most pronounced in the corpus striatum of NHP #1, which also displayed mild microhemorrhages (Figure 4). NHPs displayed mild to marked neutrophilic inflammation in the brain extending into the meninges (Figure 4). Additionally, perivascular infiltrates ranged from minimal lymphocytic, mononuclear and neutrophilic, to moderate and predominantly neutrophilic (Figure 4). The ISH staining detected substantial viral RNA in the brain tissue of all NHPs (Figure 5). The IHC staining showed mild to marked immunoreactivity of neurons in all sections of the brain with the most pronounced in the corpus striatum, thalamus, mesencephalon, and medulla oblongata (Figure 6).

**Figure 3.**
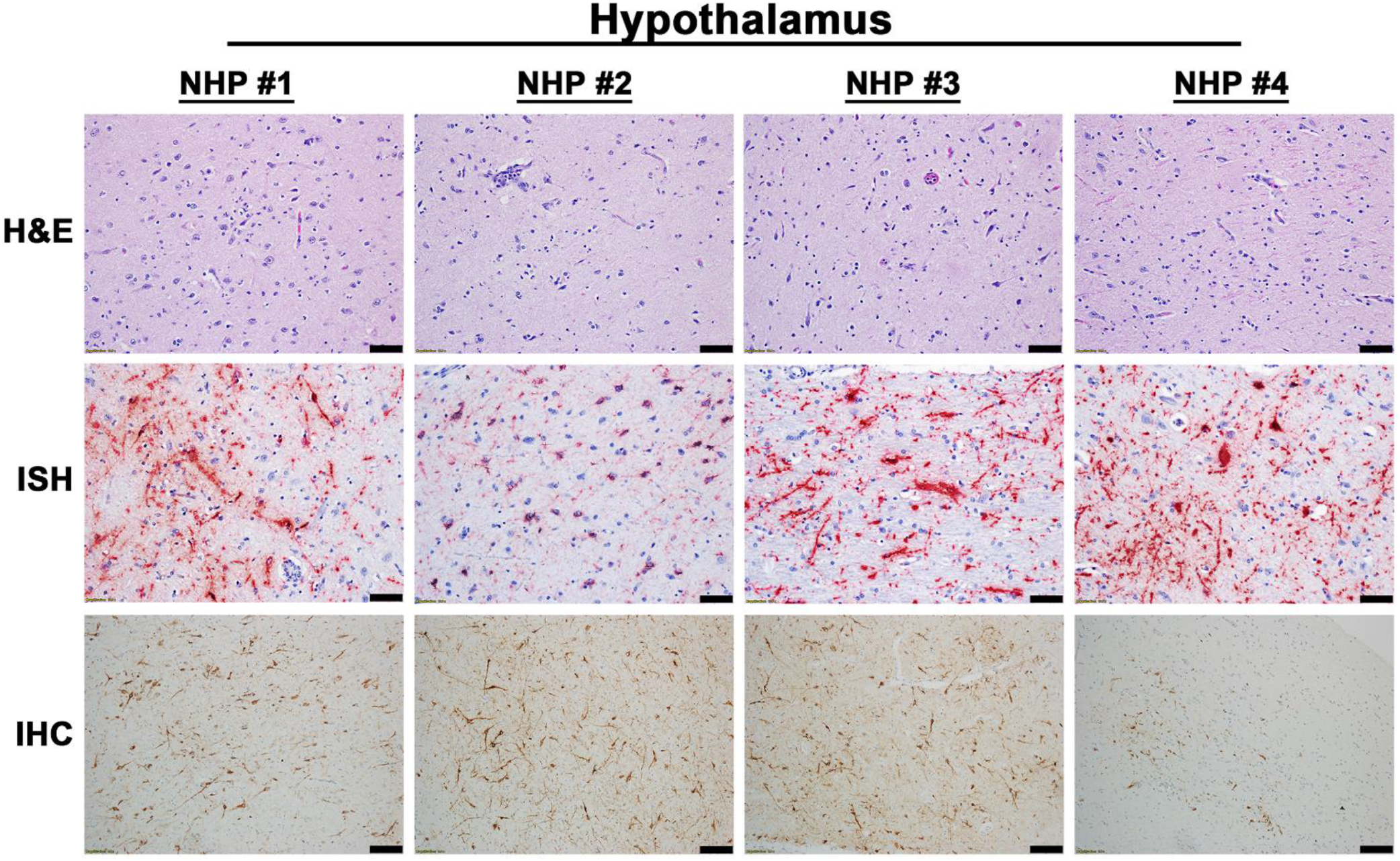
Pathology in the hypothalamus of EEEV infected cynomolgus macaques. The tissue was collected at the time of euthanasia. Hematoxylin and eosin (H&E) staining was performed to visualize histopathology. The presence of EEEV RNA and proteins was visualized via *in situ* hybridization (ISH) and immunohistochemistry (IHC), respectively. H&E, ISH, and IHC were performed on the tissues of all four NHPs. Bar = 100 um (H&E and IHC). Bar = 50 um (ISH).

**Figure 4.**
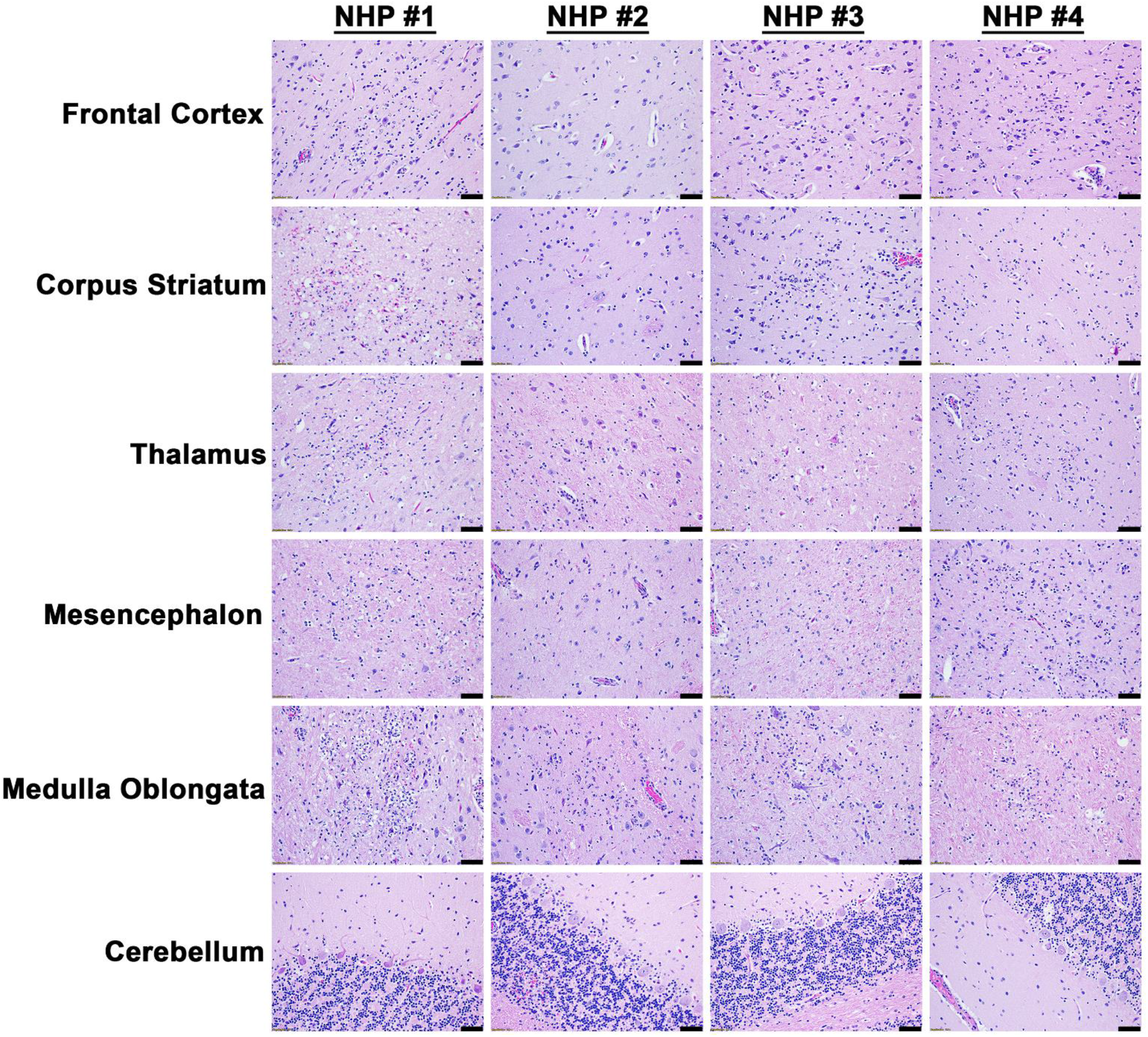
Histopathology in various parts of the brain tissues of EEEV infected cynomolgus macaques. The tissues were collected at the time of euthanasia. Hematoxylin and eosin (H&E) staining was performed to visualize histopathology. H&E was performed on the tissues of all four NHPs. Bar = 100 um.

**Figure 5.**
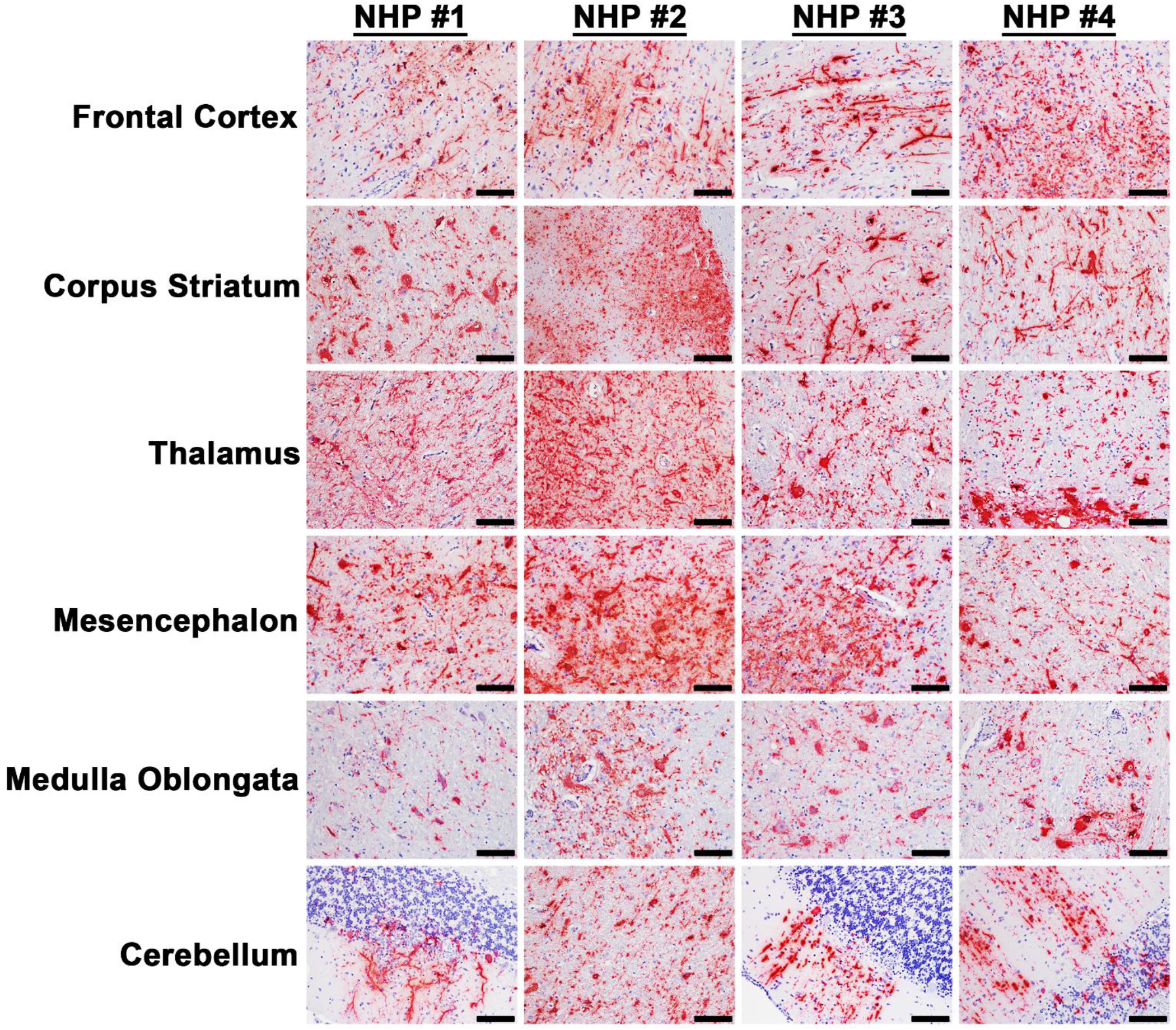
The presence of EEEV RNA in various parts of the brain tissues of infected cynomolgus macaques. The tissues were collected at the time of euthanasia. The presence of viral RNA was visualized via *in situ* hybridization (ISH). ISH was performed on the tissues of all four NHPs. Bar = 50 um.

**Figure 6.**
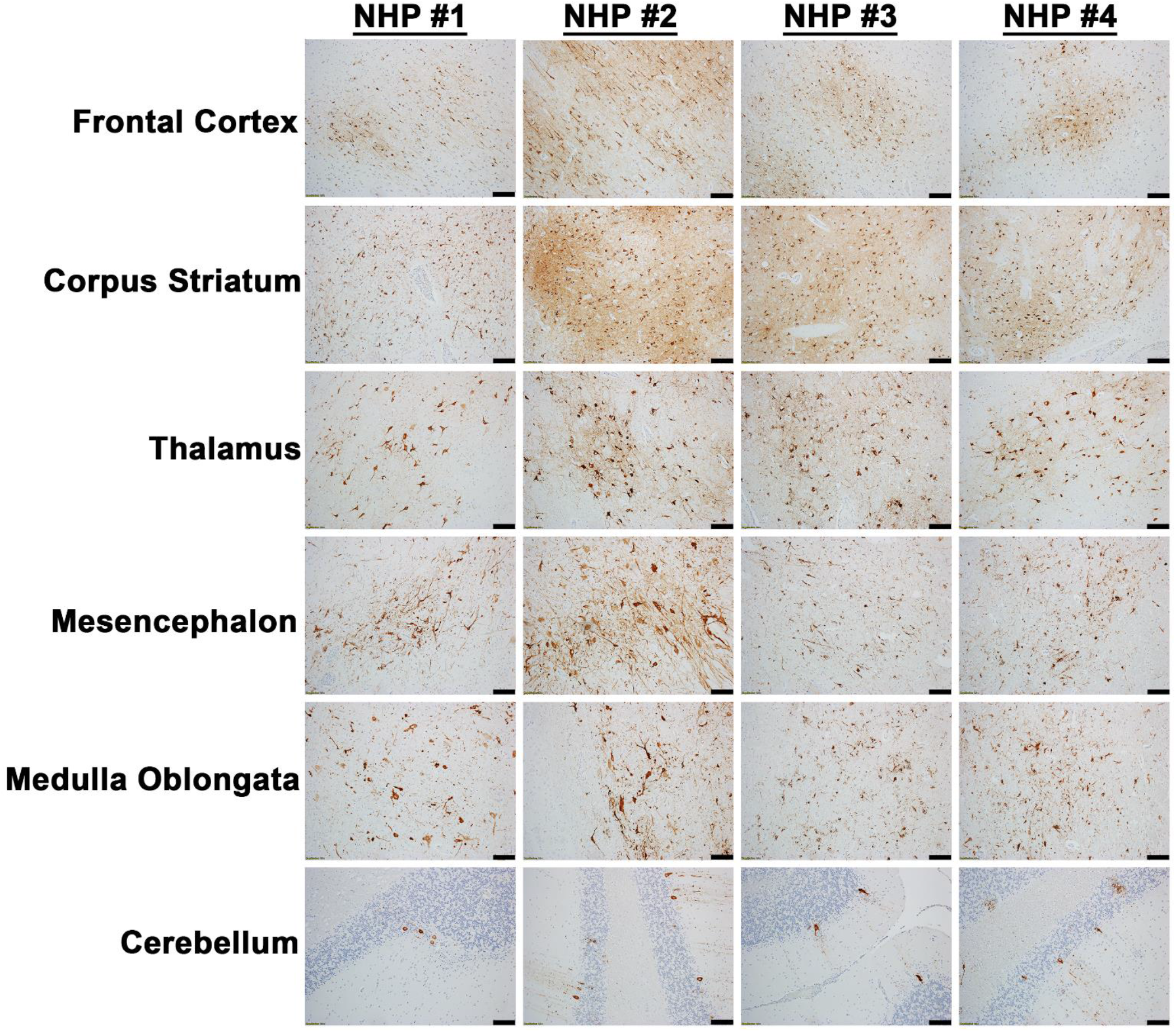
The presence of EEEV proteins in various parts of the brain tissues of infected cynomolgus macaques. The tissues were collected at the time of euthanasia. The presence of viral proteins was visualized via immunohistochemistry (IHC). IHC was performed on the tissues of all four NHPs. Bar = 100 um.

The cervical, thoracic, and lumbar spinal cord were also examined (Supp. Table 1). In contrast to the brain sections, all three sections of the spinal cord displayed minimal or no pathological lesions (Figure 7). The main feature observed in the spinal cord sections was comprised of some inflammation and myelitis. Viral RNA was readily detected in the cervical spinal cord of all four NHPs via ISH, whereas minimal or no RNA was detected in the thoracic and lumbar sections in three of the four NHPs (Figure 8). Substantial viral RNA was detected in thoracic and lumbar sections of NHP #3 (Figure 8). This finding was further verified by IHC staining that displayed a similar pattern (Figure 9).

**Figure 7.**
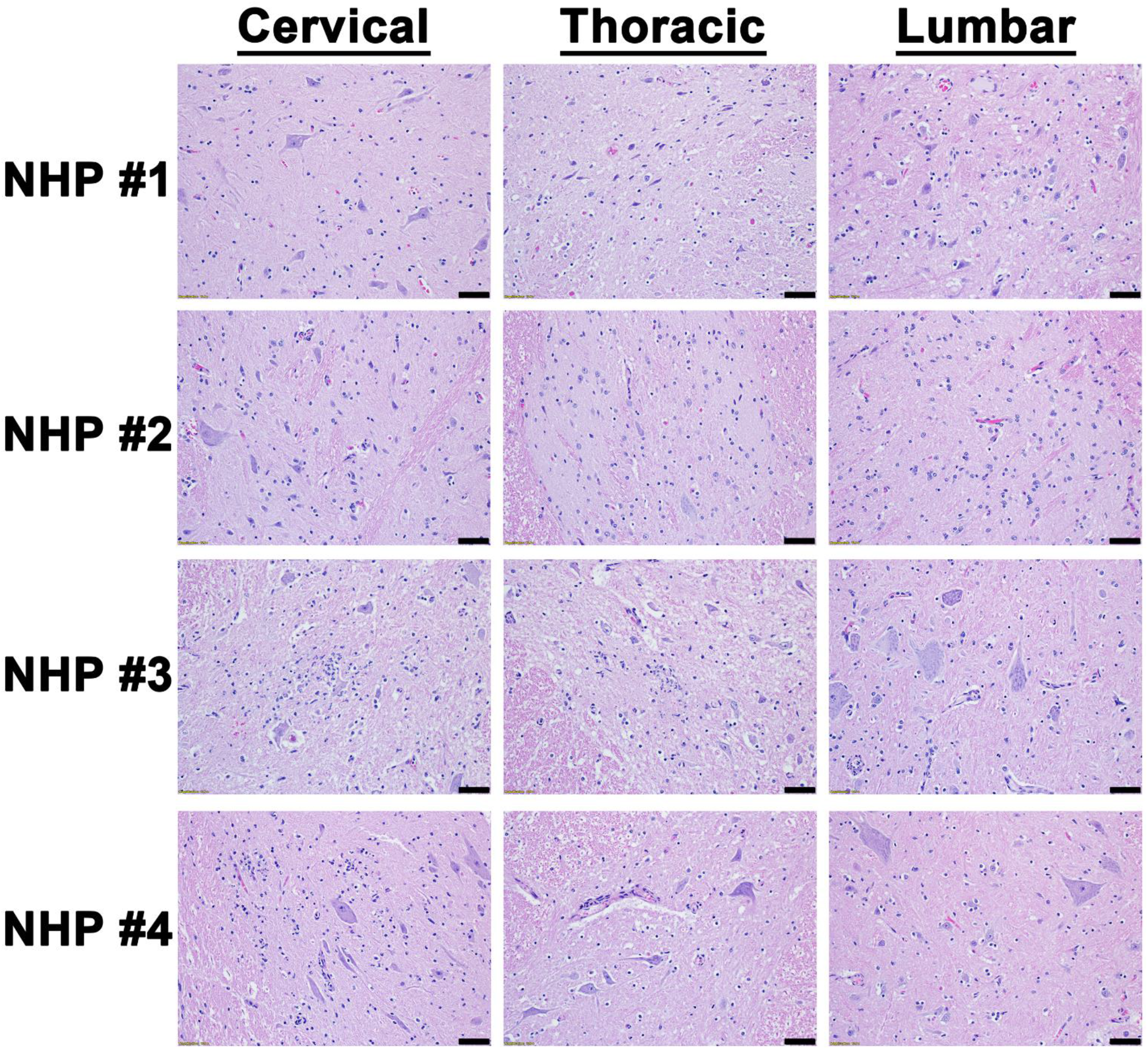
Histopathology in various parts of the spinal cord of EEEV infected cynomolgus macaques. The tissues were collected at the time of euthanasia. Hematoxylin and eosin (H&E) staining was performed to visualize histopathology. H&E was performed on the tissues of all four NHPs. Bar = 100 um.

**Figure 8.**
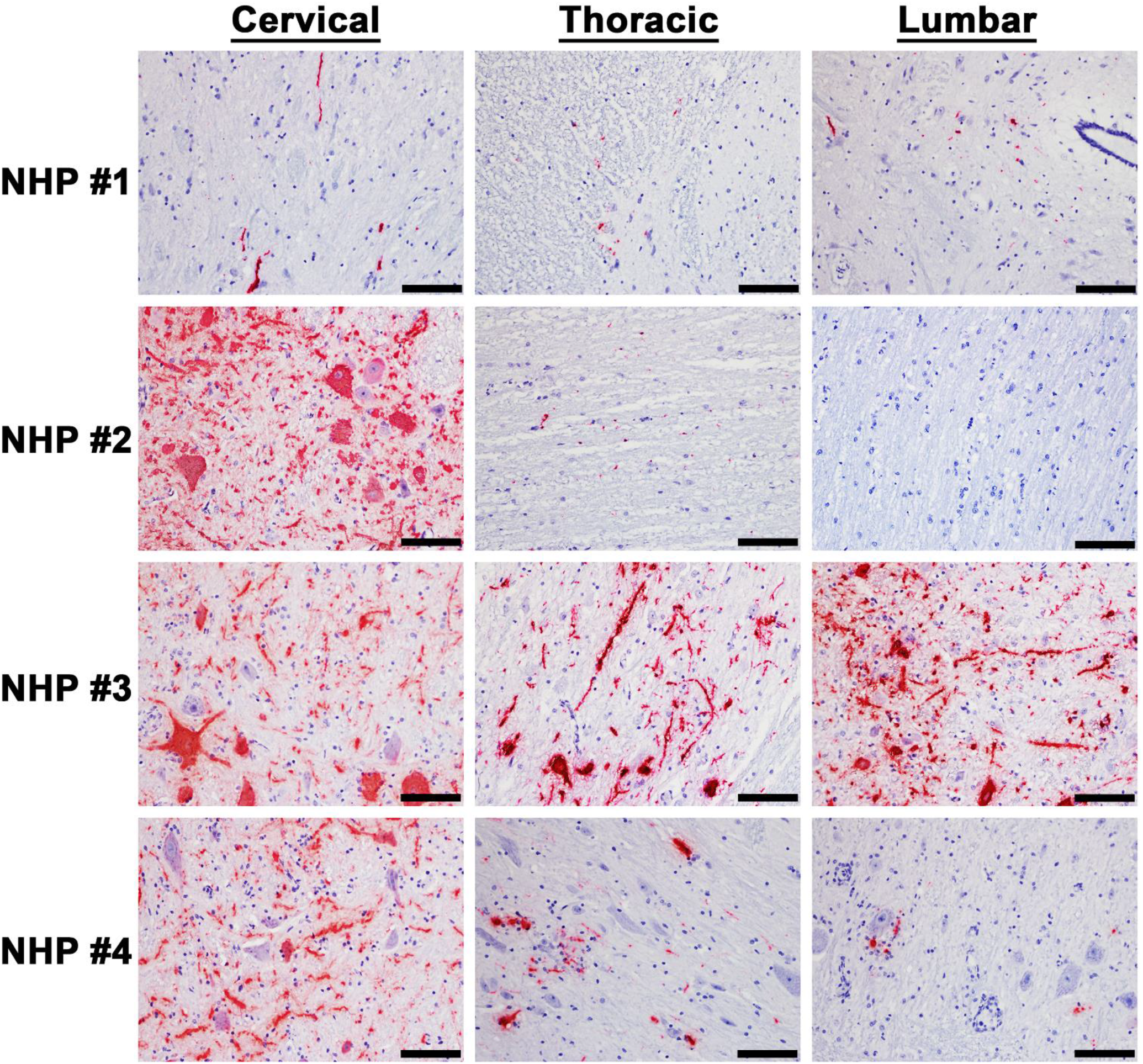
The presence of EEEV RNA in various parts of the spinal cord of infected cynomolgus macaques. The tissues were collected at the time of euthanasia. The presence of viral RNA was visualized via *in situ* hybridization (ISH). ISH was performed on the tissues of all four NHPs. Bar = 50 um.

**Figure 9.**
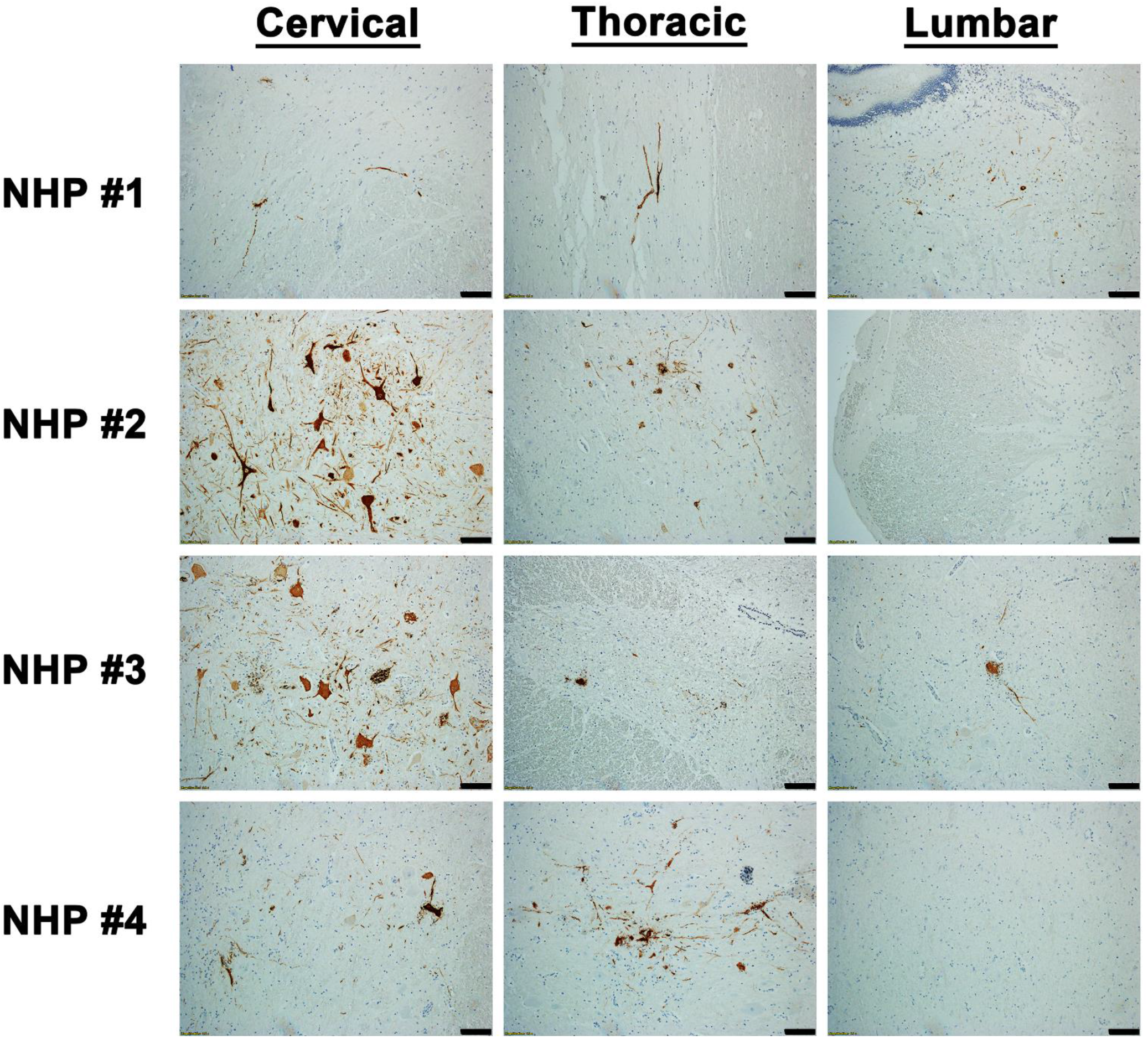
The presence of EEEV proteins in various parts of the spinal cord of infected cynomolgus macaques. The tissues were collected at the time of euthanasia. The presence of viral proteins was visualized via immunohistochemistry (IHC). IHC was performed on the tissues of all four NHPs. Bar = 100 um.

### EEEV Cell Tropism in the Thalamus of the NHPs

After establishment of EEEV infection in various brain regions, we next investigated the virus tropism in the thalamus of all infected NHPs by examine infection in the astrocytes, microglia, and the neurons. Tissue sections were stained for viral RNA and cellular markers of astrocytes (GFAP), microglia (CD68), and neurons (NeuN) (Figures 10–12). Minimal or no overlap was observed between viral RNA and GFAP or CD68, indicating minimal or no infection in the astrocytes and microglia, respectively (Figures 10 and 11). In contrast, considerable overlap of viral RNA and NeuN was observed in all NHPs suggesting that the majority of the viral infection was limited to the neurons (Figure 12).

**Figure 10.**
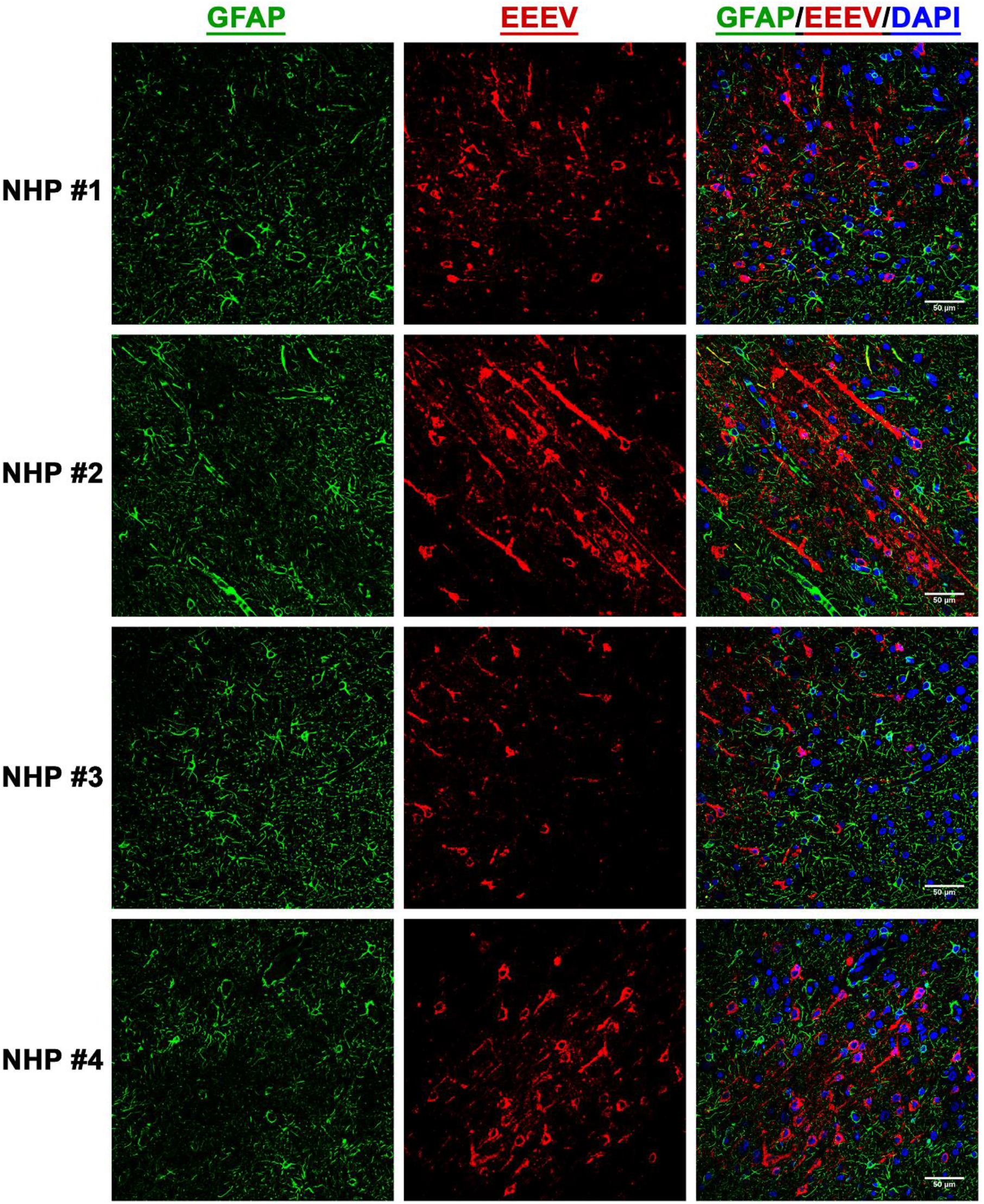
The presence of EEEV RNA in the astrocytes of infected cynomolgus macaques. Sections from the thalamus of each NHP were visualized via immunofluorescence assay. Sections were stained for GFAP (green), EEEV (red), and DAPI (blue).

**Figure 11.**
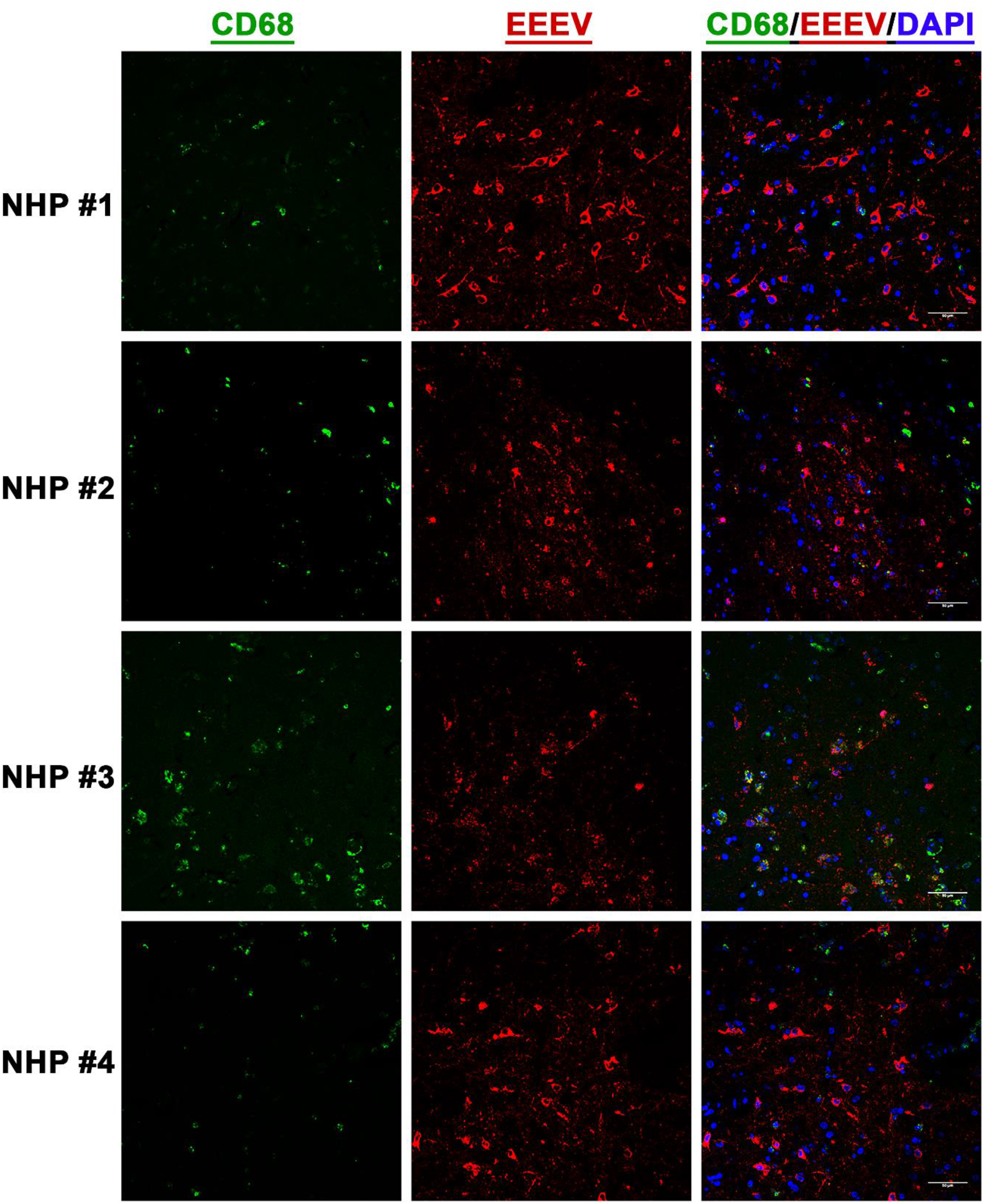
The presence of EEEV RNA in the microglia of infected cynomolgus macaques. Sections from the thalamus of each NHP were visualized via immunofluorescence assay. Sections were stained for CD68 (green), EEEV (red), and DAPI (blue).

**Figure 12.**
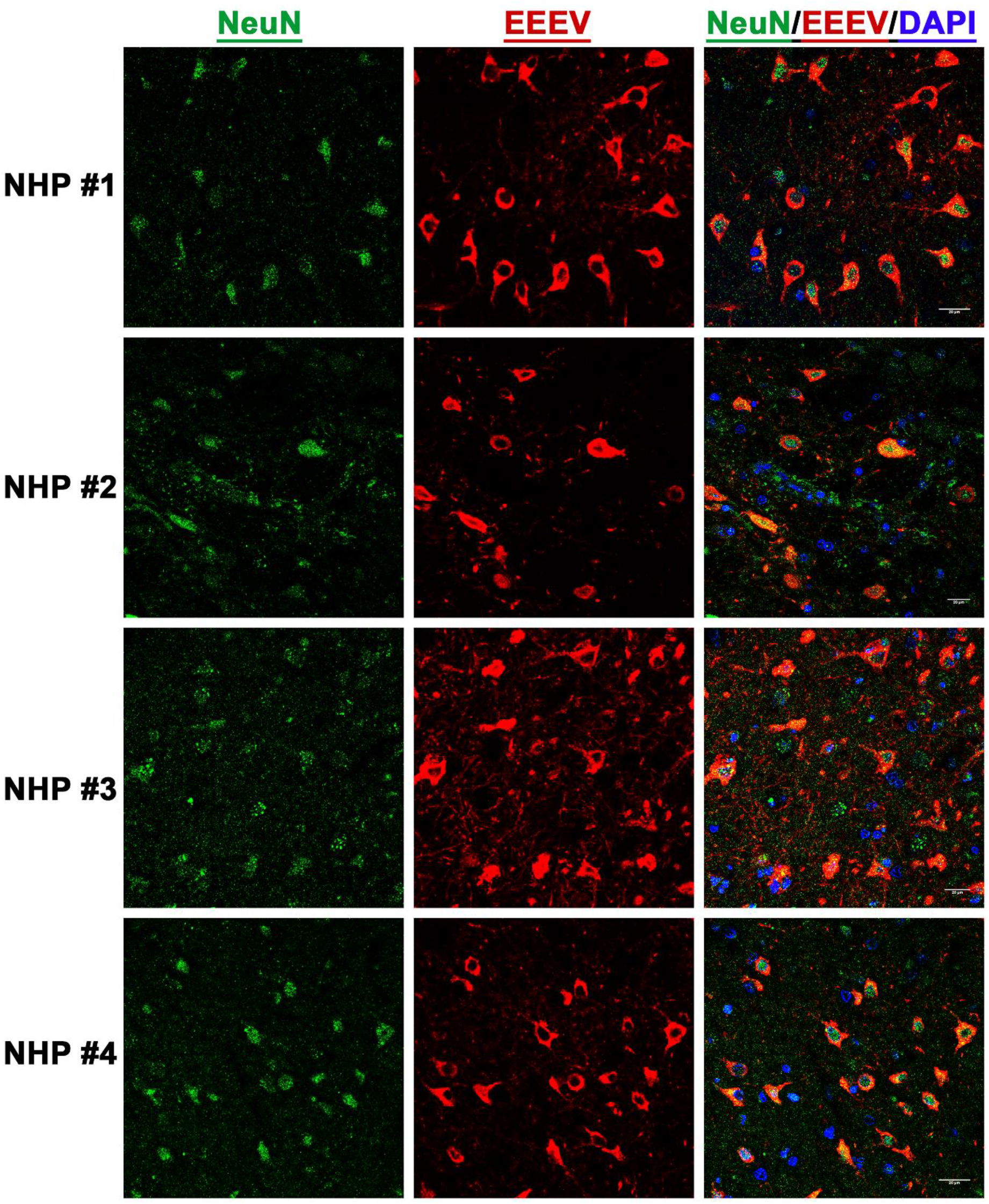
The presence of EEEV RNA in the neurons of infected cynomolgus macaques. Sections from the thalamus of each NHP were visualized via immunofluorescence assay. Sections were stained for NeuN (green), EEEV (red), and DAPI (blue).

### Localization of EEEV Virions in the Thalamus of the NHPs via Transmission Electron Microscopy (TEM)

The morphological analysis of various brain structures by the TEM showed no overt signs of apoptosis and/or necrosis as the majority of tissue sections displayed intact mitochondria and nuclei. TEM analysis showed marked presence of EEEV particles in the extracellular spaces throughout the thalamus of all NHPs (Figure 13, Supp. Figure 3). The majority of the virus particles were spherical, ~61-68 nm in diameter, and were in close proximity to plasma membranes of the surrounding cells (Figure 14). Virus particles were detected juxtaposed to myelin sheaths, surrounding the axons, as well as near synapses (Figure 15, Supp. Figure 4).

**Figure 13.**
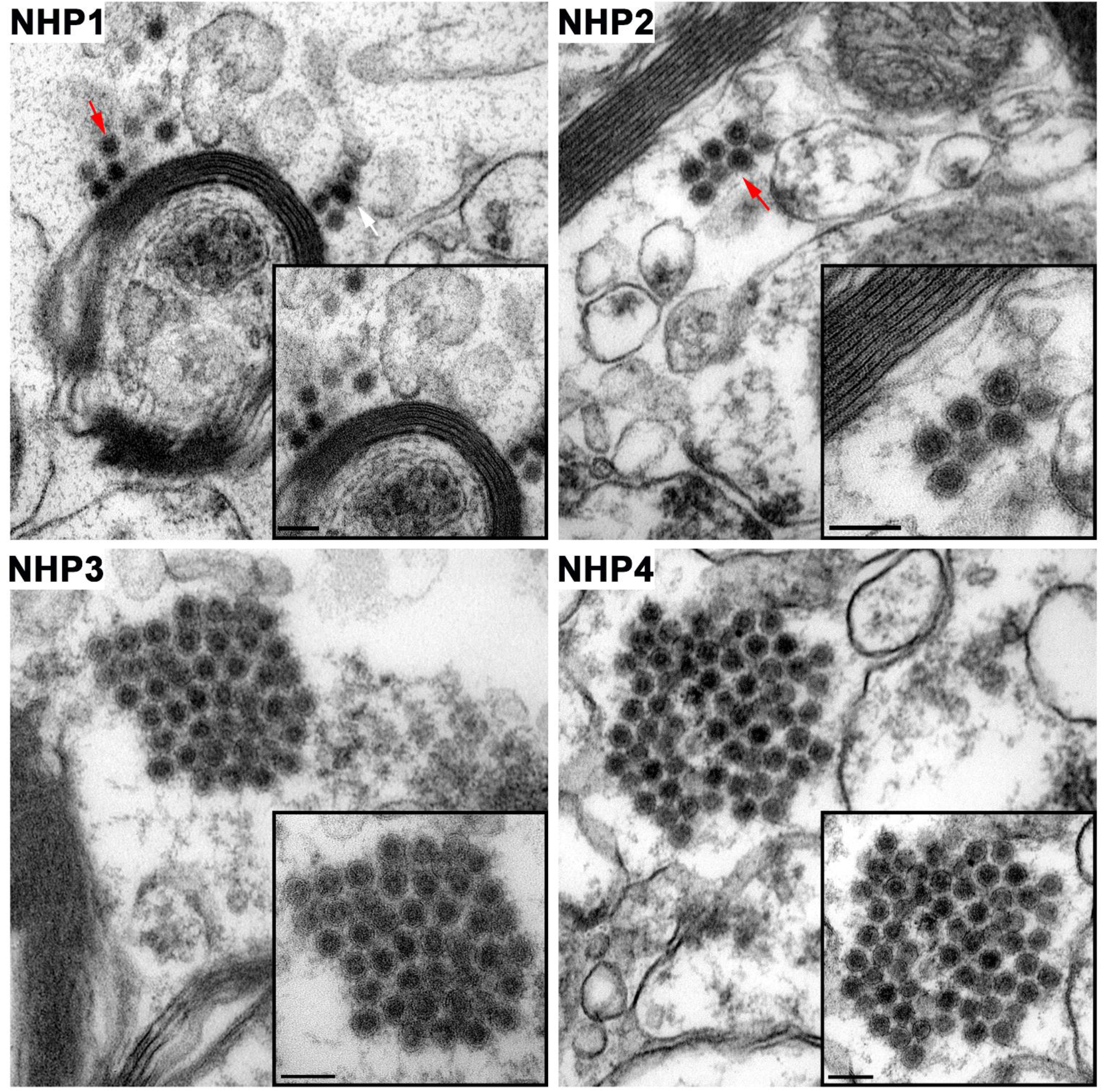
The extracellular distribution of EEEV virions in the thalamus of infected cynomolgus macaques. Sections from each NHP were examined and representative micrographs from each NHP are shown. Red arrows indicate virus particles. Bar = 200 nm.

**Figure 14.**
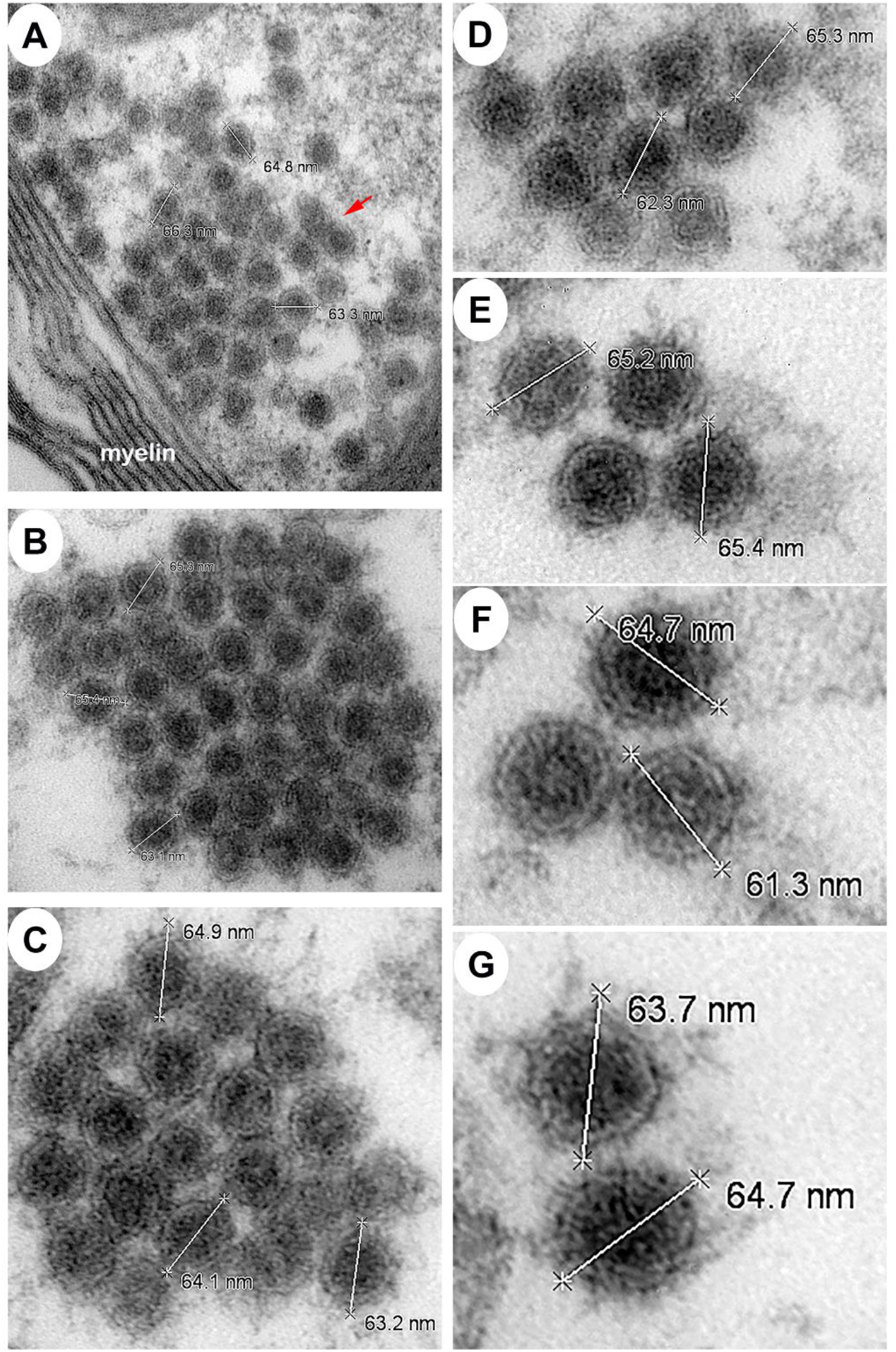
The size of extracellular EEEV virions via transmission electron microscopy (TEM). Sections from the thalamus of each NHP were examined and representative micrographs from NHPs are shown. Red arrow indicates virus particles.

**Figure 15.**
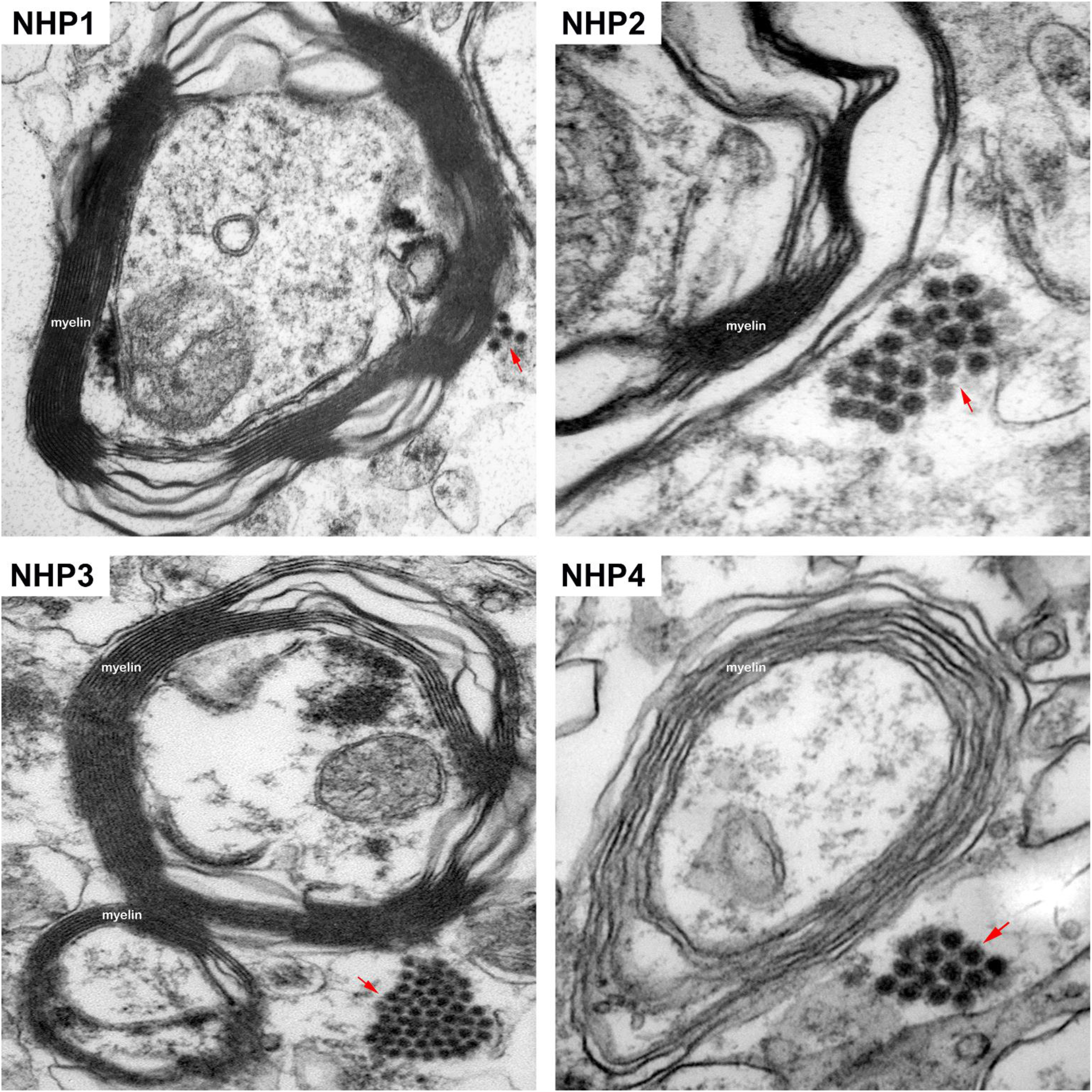
The localization of EEEV virions around the myelin sheath of neurons via transmission electron microscopy (TEM). Sections from the thalamus of each NHP were examined and representative micrographs from each NHP are shown. Red arrows indicate virus particles.

The intracellular localization of EEEV within the thalamus was examined by detecting the presence of EEEV particles within the axons of neurons. Virus particles, ~62-67 nm in diameter, were detected within the axons in all NHPs (Figure 16). Surprisingly, the majority of the particles were not contained within vesicles and appeared to be free virions. In two sequential sections, ~80 nm apart, of an axon, the quantity of EEEV virions present inside an axon was assessed. The sections showed the presence of 18 and 17 particles, respectively (Figure 17 A and B). This finding highlights the potential of large quantity of particles that can migrate through a single axon to infect other neurons.

**Figure 16.**
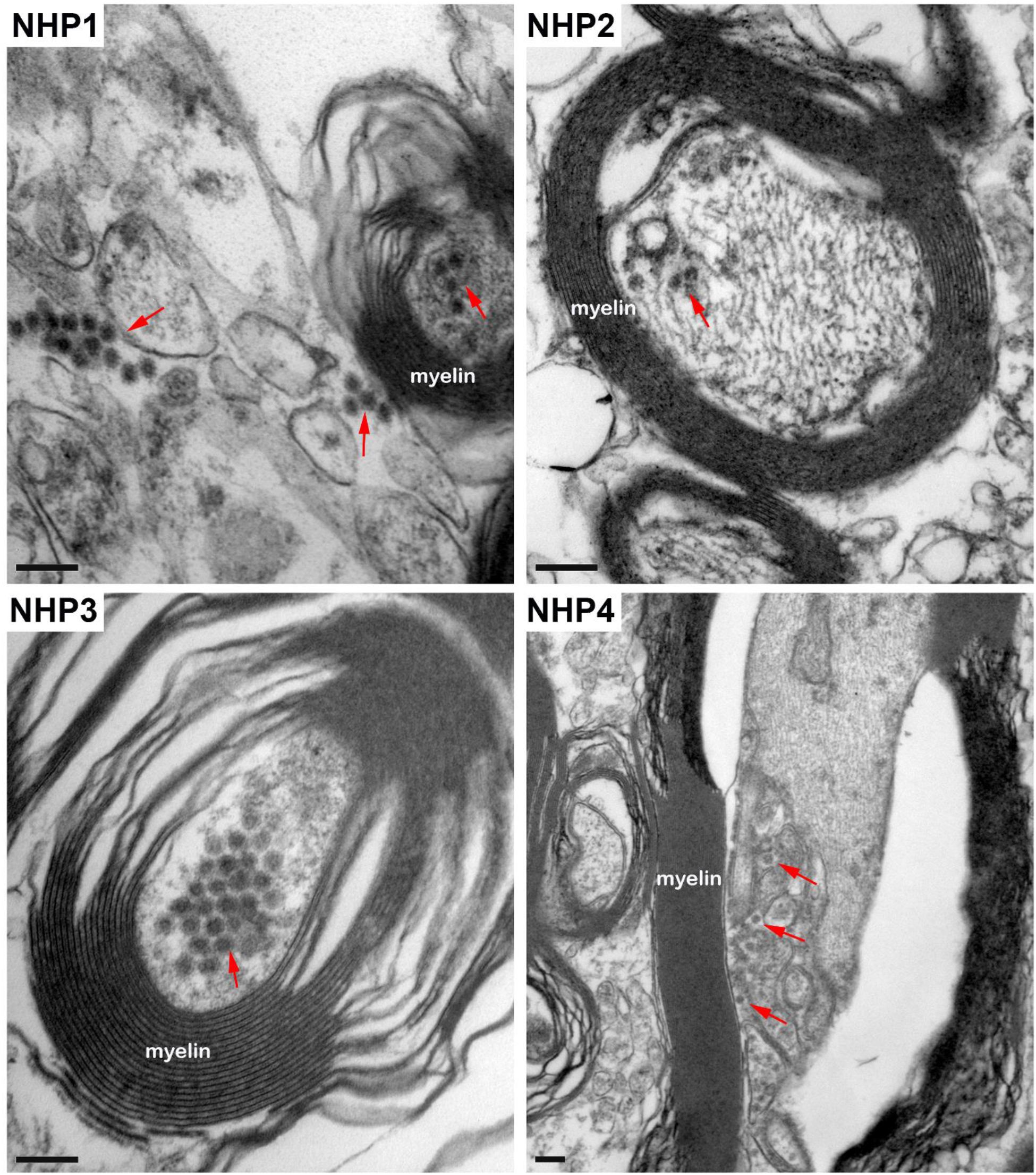
The localization of EEEV virions within the axons of neurons via transmission electron microscopy (TEM). Sections from the thalamus of each NHP were examined and representative micrographs of each NHP are shown. Red arrows highlight virus particles. Bar = 200 nm.

**Figure 17.**
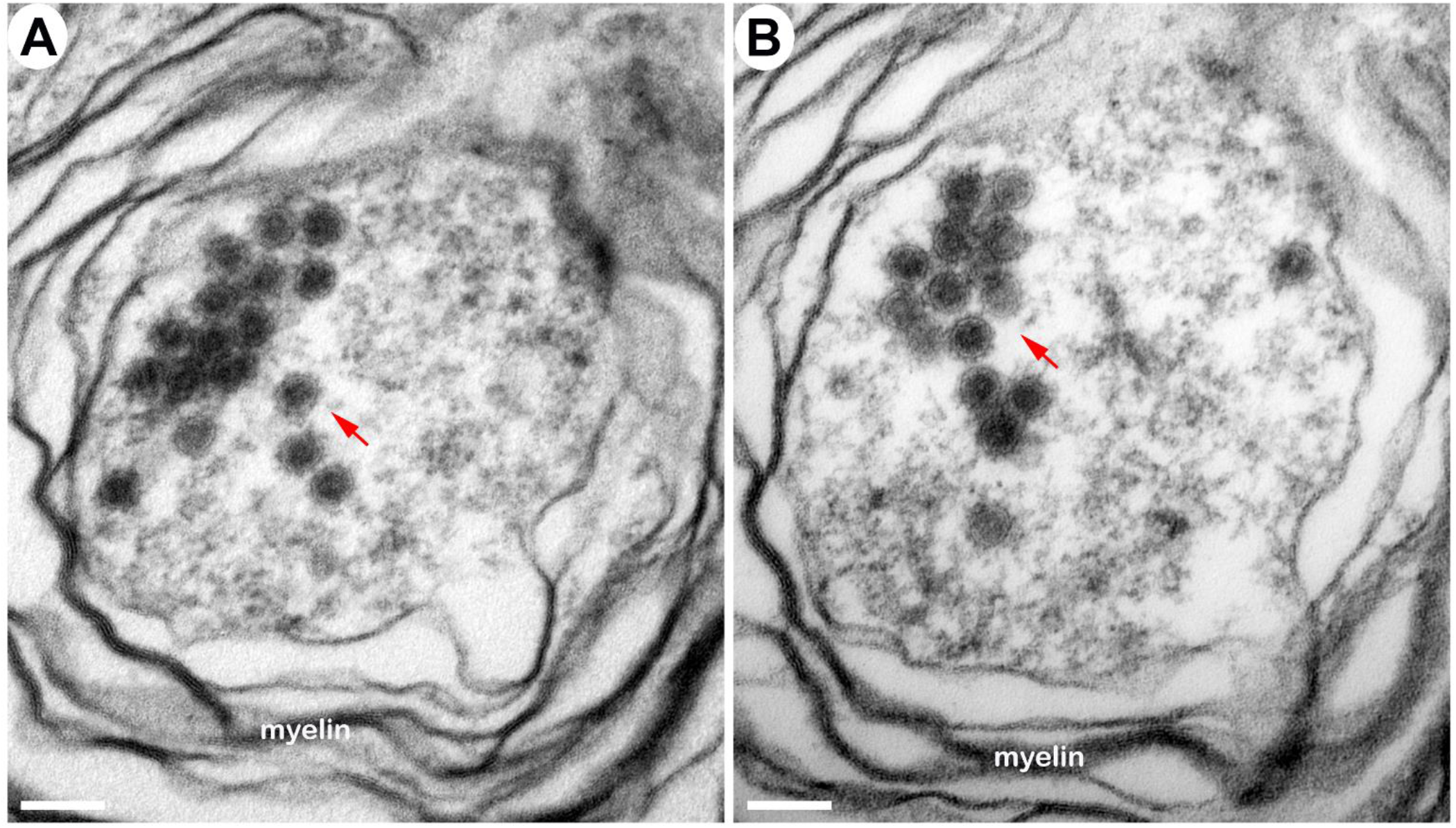
The localization of EEEV virions within an axon of a neuron via transmission electron microscopy (TEM). Sections from the thalamus of NHP #3 were examined. Two sequential sections, ~80 nm apart, of an axon are shown. Red arrows indicate virus particles. Scale bar = 100 nm.

Next, we sought to investigate active virus replication centers by detecting cytopathic vacuoles and budding virions in infected cells. Cells with active replication were rare in the brain, however, they were detected in all NHPs. The majority of the virus replication was localized to the amygdala, hippocampus, thalamus, and hypothalamus (Figures 18 and 19, Supp. Figure 5). Extensive cytopathic vacuoles with attached and free nucleocapsid, ~40 nm in diameter, were present within the cytoplasm of infected cells (Figures 18 and 19). Furthermore, infectious virions ~65-68 nm in diameter were observed budding from infected host cell plasma membrane (Figure 19).

**Figure 18.**
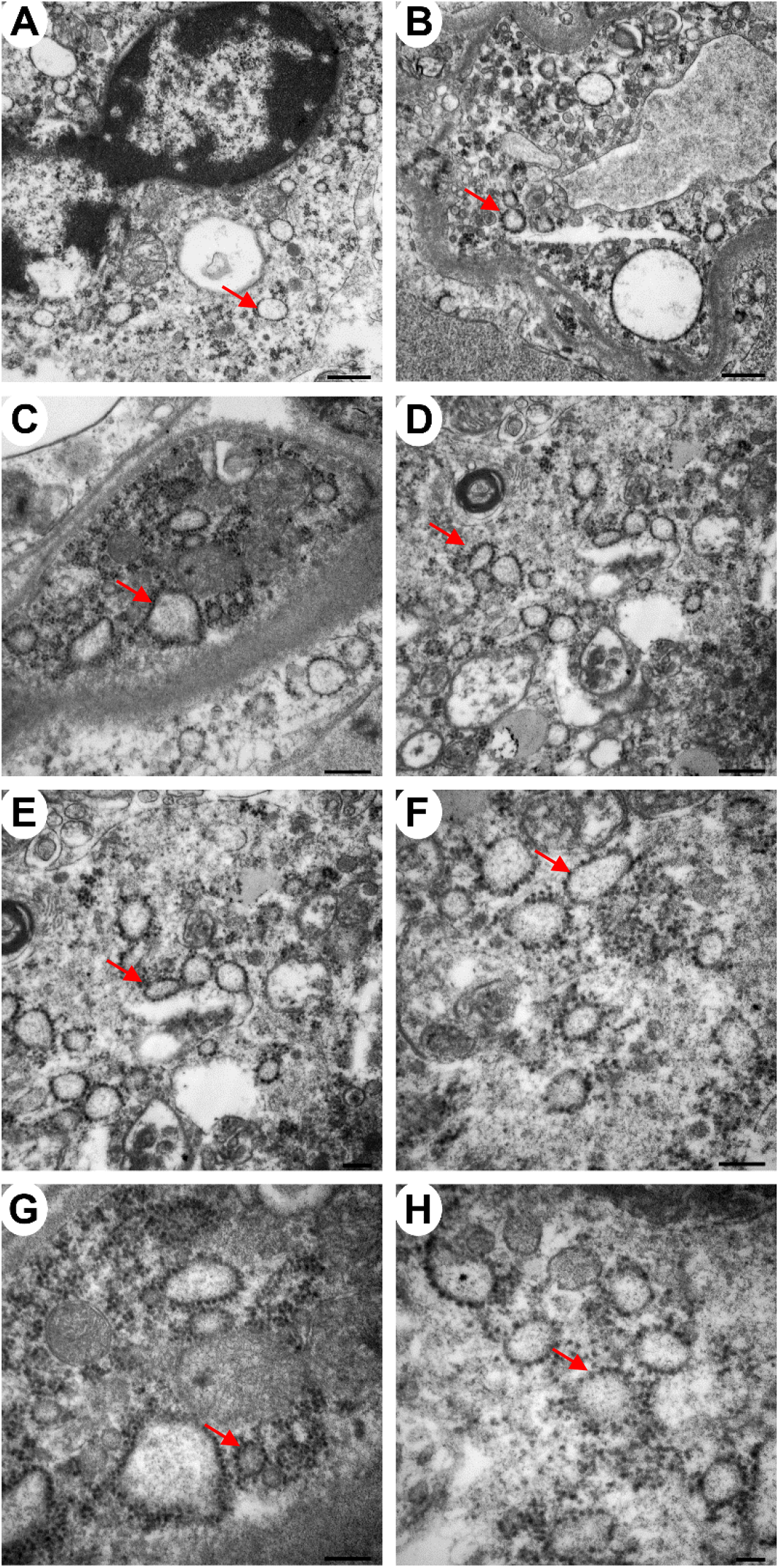
The detection of cytopathic vacuoles in the cytoplasm of EEEV infected cells via transmission electron microscopy (TEM). Sections from the thalamus of infected NHPs were examined. Micrographs of NHP #4 are shown. Scale bars: Panels A and B = 600 nm, C and D = 400 nm, E, F and G = 200 nm, and H =100 nm.

**Figure 19.**
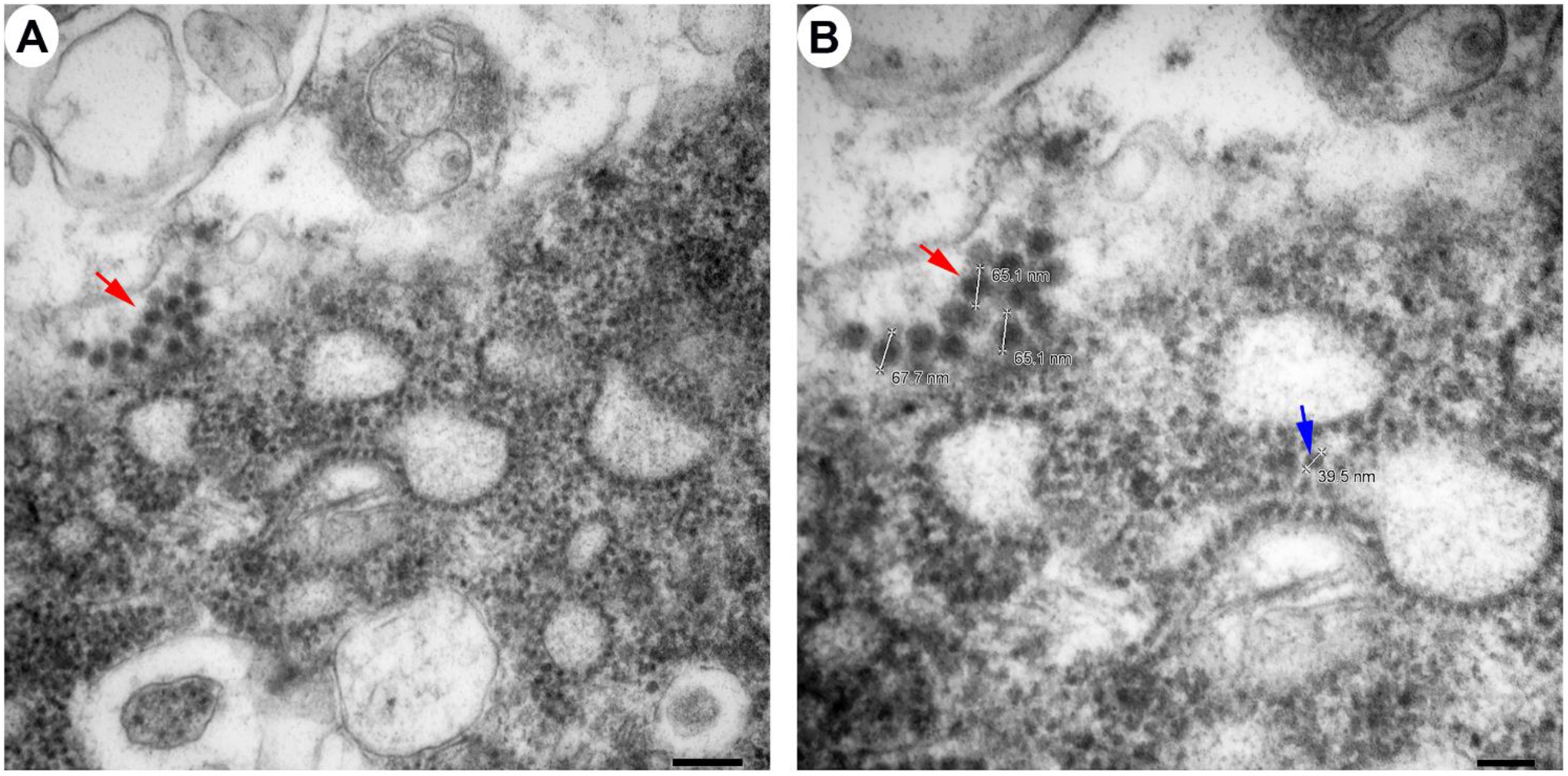
The detection of cytopathic vacuoles, nucleocapsid, and budding virions in EEEV infected cells via transmission electron microscopy (TEM). Blue and red arrows indicate nucleocapsid and virus particles, respectively. Sections from the thalamus of infected NHP #3 were examined. Scale bars: A = 200 nm, B = 100 nm.

Another interesting finding in the TEM experiments was the presence of virus particles enclosed within undefined vesicular compartments in the extracellular space of all NHP tissues (Figure 20). The vesicles were composed either exclusively of virions or mixture of particles and cellular component of similar size and shape (Figure 20). Free virions could also be found adjacent to the enclosed vesicle (Figure 20 A and C).

**Figure 20.**
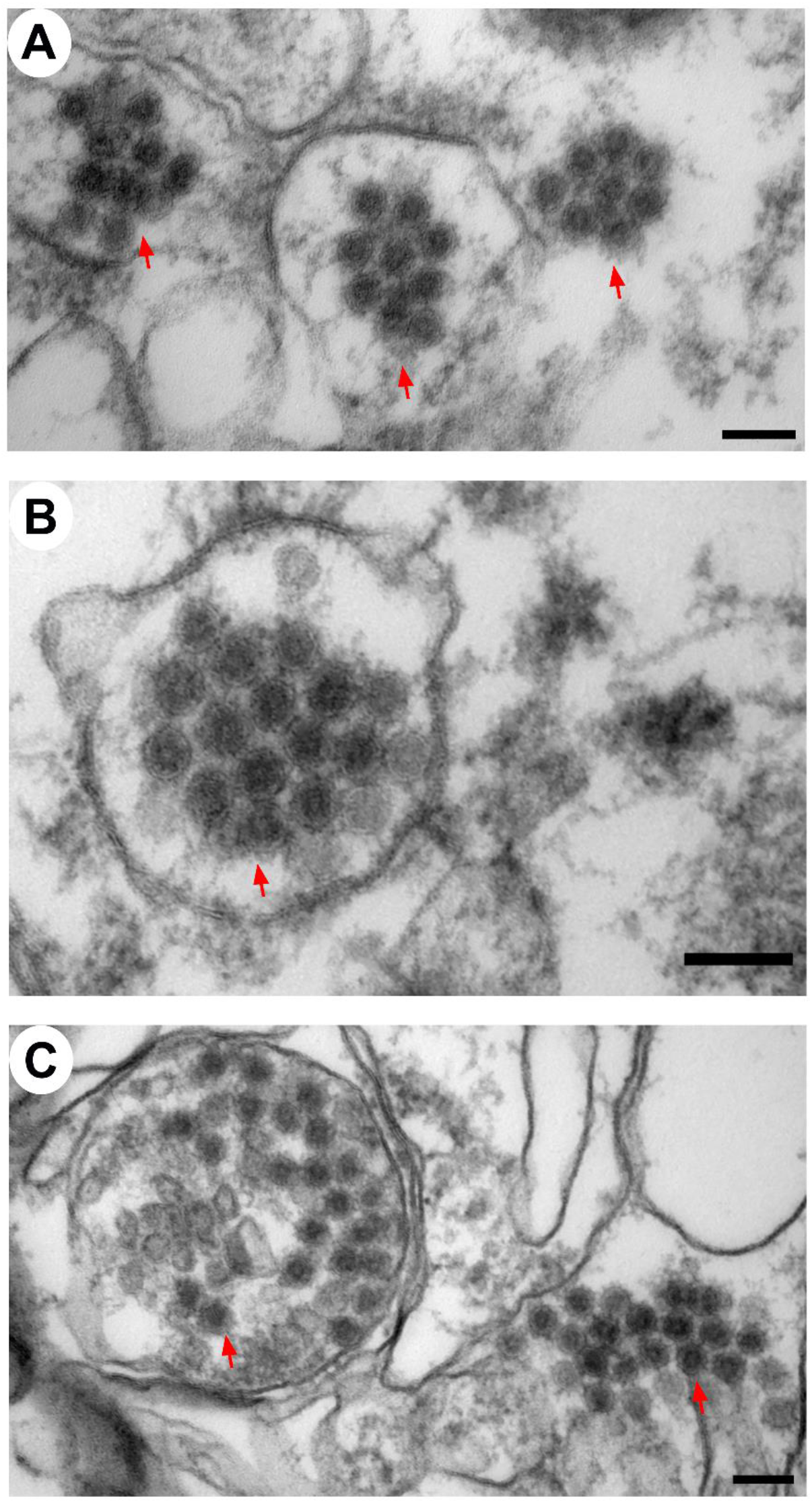
The detection of EEEV particles enclosed within vesicles via transmission electron microscopy (TEM). Sections from the thalamus of infected NHPs were examined and representative micrographs are shown. Red arrows indicate virus particles. Scale bar = 100 nm.

Although rare, necrotic lesions were visible within the thalamus and were detected by TEM. Considerable degeneration of the cellular architecture was observed with loss membrane integrity, disintegration of organelles, and cell lysis (Figure 21). EEEV particles were readily detected scattered throughout the remaining cell debris (Figure 21).

**Figure 21.**
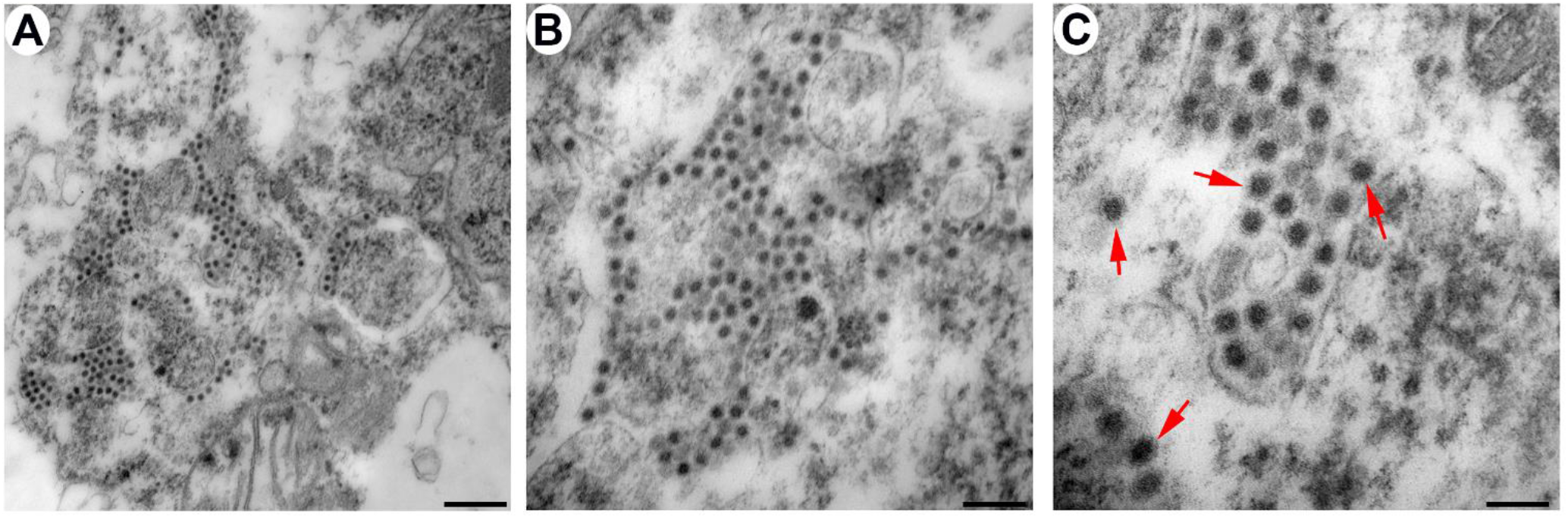
The detection of necrotic lesions in the thalamus of NHP #1 via transmission electron microscopy (TEM). Red arrows indicate virus particles. Scale bars: A = 400 nm, B = 200 nm, C = 100 nm.

## DISCUSSION

The susceptibility of cynomolgus macaques to North American lineage of EEEV via the aerosol route has been explored previously [8–10]. However, indepth pathology studies have not been performed to gain insights into the mode of virus dissemination following aerosol challenge. Following challenge, there are two potential routes of virus dissemination in the NHP host. The initial virus replication in the respiratory tract followed by systemic infection and subsequent access to central nervous system (CNS). Alternatively, the olfactory epithelium and bulb could serve as the initial site of virus replication followed by virus transport and infection of the olfactory tract with spread to the distal regions of the brain. The data from our study showed no evidence of gross and/or microscopic changes, as well as viral RNA or proteins in the lung, liver, heart, spleen, and kidneys. In contrast, pathological lesions were detected in the brain comprising of neutrophilic inflammation, neuronal degeneration, and necrosis. EEEV RNA and proteins were also readily detected throughout many parts of the brain and spinal cord. A gradation of viral RNA and proteins was observed the most in the cervical region due to its proximity to the brainstem and least in the lumbar region. Lastly, the data from our companion manuscript showed presence of high EEEV infectious titers at the olfactory bulb of all NHPs. Taken together, these data support the rapid and direct spread of EEEV via the olfactory bulb into the brain followed by dissemination into the spinal cord.

The dissemination of EEEV following aerosol infection has not been investigated in previous macaque studies; however, it has been examined in mice, guinea pigs, and marmosets [11–14]. In these previous studies, virus was localized almost exclusively in the brain and was readily detected in the frontal cortex, corpus striatum, thalamus, hippocampus, mesencephalon, pons, medulla oblongata, and cerebellum [11–14]. In contrast, EEEV could not be detected in the heart, liver, lung, spleen, and kidney of guinea pigs and marmosets [11, 14]. Murine studies displayed similar pattern to guinea pigs and marmosets, however, EEEV was detected lung and heart [12, 13]. Our NHP data are in agreement with the guinea pig and marmoset studies. The presence of virus in the mouse lung and heart tissues shows important differences between the murine and other animal models.

In nature, EEEV is transmitted via a mosquito bite and can cause fatal encephalitis in many mammalian species including horses, sheep, cattle, alpacas, llamas, deer, dogs, pigs, and humans [15–57]. Following the bite of an infected mosquito, the virus replicates locally in skeletal muscle cells, fibroblasts, and osteoblasts, gains access to the peripheral tissues and organs, and eventually disseminates into the CNS to cause fatal encephalitis [58]. During the course of infection, extensive pathology is observed in the visceral tissues and organs including lungs, liver, kidneys, spleen, intestine, as well as cardiac and skeletal muscle [21, 23, 26, 28, 48, 52, 57]. The pathology is comprised of severe pulmonary edema and congestion, multifocal hemorrhage, splenic atrophy, myocarditis, and necrosis [21, 23, 26, 28, 48, 52, 57]. The lack of similar pathology in visceral organs and tissues in animals infected via an aerosol route demonstrates that the route of infection substantially alters the virus dissemination [11–14].

EEEV localizes in the CNS of many mammalian species including humans regardless of the route of infection [9–30, 32–47, 49–66]. Virus can be readily detected in basal ganglia, hippocampus, frontal cortex, pons, thalamus, substantia nigra, mesencephalon, medulla oblongata, cerebellum, and spinal cord with minimal to moderate lesions [9–30, 32–47, 49–66]. These microscopic findings consist of neuronal degeneration and necrosis, neuropil vacuolation, gliosis, and satellitosis, neuronophagia, lymphocytic perivascular cuffing, lymphocytic meningitis, perivascular cuffs, neutrophil infiltrate, and microhemorrhage. The tropism of EEEV is predominantly limited to the neurons, however, astrocytes and microglia cells are also infected. The results of our study are in agreement with the majority of previously reported findings, however, there are several important differences. First, in our study, the majority of the cellular architecture in all brain regions remained intact and the focal degenerative and necrotic lesions were limited to the amygdala, hippocampus, corpus striatum, thalamus, mesencephalon, and medulla oblongata. Second, neutrophils comprised the majority of inflammatory infiltrates, whereas minimal lymphocytic infiltrates were observed. Third, the tropism of EEEV was almost exclusively to neurons. Fourth, microscopic findings were either absent or minimal in all sections of the spinal cord. These differences highlight that aerosol infection can substantially alter virus pathogenesis.

Limited studies have examined EEEV pathogenesis in the brain utilizing TEM [16, 29, 36, 67]. These studies showed the presence of infectious particles, ~55-60 nm in diameter, localized almost exclusively in the extracellular spaces. The evidence of virus replication was either absent or rare in the tissues. Cytopathic vacuoles and nucleocapsid, ~28 nm in diameter, were observed in the cytoplasm of infected neurons and microglia. Infected and uninfected neurons, astrocytes, and microglia displayed dilated rough endoplasmic reticulum. Our study is in agreement with most of the previously reported findings, with one exception with regards to the size of virus particles and nucleocapsid. The infectious particle and nucleocapsid size was smaller in the previous human and mouse TEM studies than the recent cryo-electron microscopy (cryo-EM) studies that estimate the infectious particle and nucleocapsid size of ~65-70 and ~40-45 nm, respectively [3, 68–70]. Our study is in agreement with the latter data. One potential explanation for this discrepancy is the shrinking effects of formalin fixation, dehydration, and paraffin embedding. The process of inactivation and embedding can reduce tissue size by up to 15% [71–74].

Axonal transport is an essential homeostatic process responsible for movement of RNA, proteins, and organelles within neurons [75]. Viruses including rabies, polio, West Nile, and Saint Louis encephalitis can utilize this critical mechanism and disseminate in the CNS via neuron-to-neuron spread [76–78]. The data from the present study showed that viral replication was limited to the olfactory bulb and proximal regions including the amygdala, hippocampus, thalamus, and hypothalamus. Viral RNA, proteins, and infectious particles were also detected in distal parts of the brain, however, minimal or no virus replication was detected. In addition, the infectious particles were present in the axon of neurons in all four NHPs. Thirty-five infectious particles were observed in a single 160 nm section of an axon. Taken together, these data strongly suggest that EEEV is able to rapidly spread throughout the CNS following aerosol challenge likely via axonal transport and warrants further investigation.

Many of the physiological parameters measured with advanced telemetry and 24-hr continuous monitoring were considerably altered following infection; temperature +3.0-4.2 °C, respiration rate +56-128%, activity −15-76%, +5-22%, heart rate +67-190%, systolic blood pressure +44-67%, diastolic blood pressure +45-80%, ECG abnormalities, reduction in food/fluid intake and sleep, and EEG waves −99-+6,800%. Many of these parameters are under the control of the autonomic nervous system (ANS). The master regulator of the ANS is the hypothalamus which is comprised of numerous important nuclei that regulate these parameters; preoptic area (temperature), suprachiasmatic nuclei (circadian rhythm), paraventricular nuclei and supraoptic nucleus (hunger/satiety), tuberomamillary nucleus and the perifornical lateral (sleep), arcuate nucleus and paraventricular nucleus (blood pressure), arcuate nucleus (cardiac electrical system and heart rate), paraventricular nucleus, perifornical area, and dorsomedial hypothalamus (respiration) [79–86]. The hypothalamic nuclei are interconnected with many other regulatory centers such as the thalamus, basal ganglia, medulla oblongata, and others to exert control on important physiological parameters. The histopathology, ISH, IHC, and TEM data from this study shows the presence of viral RNA, proteins, and replication centers in the ANS control centers. These data suggest that EEEV infection in the brain likely produces disruption and/or dysregulation of the ANS control centers to produce rapid and extreme alterations in physiology and behavior to cause severe disease.

In many regions of the brain, EEEV infection produced minimal necrosis and inflammatory infiltrates, and majority of the cellular architecture remained intact. Accordingly, the neuronal necrosis and/or host inflammation cannot alone explain the fatal disease in the NHPs. One potential explanation of these results is that EEEV pathogenesis, in part, may be due to rapid local and global neuronal dysfunction. This hypothesis has been investigated for a prototypic encephalitic virus, rabies virus (RABV). RABV can exert neuronal dysfunction by multiple mechanisms. RABV infection in neurons can induce degeneration of axons and dendrites without inflammation or cell death, axonal swelling, generation of toxic metabolites such as reactive oxygen species, decreased expression of housekeeping genes, impairment of both the release and binding of serotonin, and reduction in expression of voltage-dependent sodium channels [87–94]. Similar to RABV, the axonal transport of EEEV may also disrupt the transport of RNA, proteins, and/or organelles to produce neuronal dysfunction and leading to fatal outcomes. This hypothesis requires further investigation to elucidate the potential mechanism/s.

As described in our companion manuscript, NHP #1 experienced a critical cardiovascular event and was subsequently euthanized. The investigation of the cardiac tissue showed no evidence of viral induced pathology, RNA, proteins, or host inflammatory response. In contrast, the brain tissues displayed some microscopic lesions as well as considerable presence of viral RNA and proteins, particularly in the hypothalamus and medulla oblongata. These data suggest EEEV infection of the ANS control centers may have led to the dysregulation and/or disruption of the heart’s electrical activity leading to a critical cardiac event. Lastly, the electrolyte imbalance due to considerable decrease in food/fluid intake in the NHP prior to the cardiac event may also contribute to the disruption and dysfunction the heart’s electrical activity.

There are several important implications of our EEEV study in regard to countermeasure development. First, the exposure by the aerosol route produces a rapid and profound infection of the CNS including the ANS control centers. Second, the axonal transport likely facilitates substantial neuron-to-neuron spread of virus. Third, the rapid viral spread in the CNS leads to considerable alterations of critical physiological parameters as early as ~12-36 hpi suggesting that the post-aerosol challenge window for therapeutic intervention may be very short in the NHP model. Fourth, the presence of infectious virus within axons and the subsequent potential spread via axonal transport demonstrate the necessity for targeting small molecule or antibody therapeutics inside the axons to prevent/reduce infection and transport. Fifth, the investigation of therapeutics and vaccines in an aerosol NHP model should include monitoring of brain waves and comprehensive brain pathology following challenge.

In summary, NHPs exhibited considerable alternation in many important physiological parameters within early as ~12 hpi following EEEV aerosol challenge. EEEV initially replicated at olfactory bulb and was rapidly transported to distal parts of the brain likely utilizing axonal transport to facilitate neuron-to-neuron spread. Once within the CNS, the virus infected the ANS control centers to likely cause the disruption and/or dysregulation of critical physiological parameters leading to NHPs meeting the euthanasia criteria ~106-140 hpi. The lack of diffuse necrosis in the CNS and ANS suggests that EEEV pathogenesis is in part likely due to neuronal dysfunction and is an important parameter for the evaluation of countermeasure development.

## MATERIALS AND METHODS

### Virus

Eastern equine encephalitis virus isolate V105-00210 was obtained from internal USAMRIID collection. The details of the stock are described in our companion manuscript. Briefly, the virus stock was deep sequenced to verify genomic sequence and to ensure purity. In addition, the stock was tested to exclude presence of endotoxin and mycoplasma.

### Ethics Statement

This work was supported by an approved USAMRIID IACUC animal research protocol. Research was conducted under an IACUC approved protocol in compliance with the Animal Welfare Act, PHS Policy, and other Federal statutes and regulations relating to animals and experiments involving animals. The facility where this research was conducted is accredited by the Association for Assessment and Accreditation of Laboratory Animal Care, International and adheres to principles stated in the Guide for the Care and Use of Laboratory Animals, National Research Council, 2011 [95].

### Non-human Primate Study Design

Study design is detailed in our companion manuscript. Briefly, four [2 males (NHPs #1 and 4), 2 females (NHPs #2 and 3)] cynomolgus macaques (*Macaca fascicularis*) of Chinese origin were obtained from Covance and were challenged with a target dose of 7.0 log_10_ PFU of EEEV via the aerosol route. Following challenge, all four NHPs exhibited severe disease and met the euthanasia criteria ~106-140 hpi. Lung, liver, spleen, kidney, heart, spinal cord, and brain tissues were collected from each NHP at the time of euthanasia. Tissues were fixed for >21 days in 10% neutral buffered formalin.

### Tissues Processing and Histopathology

Tissue sections from various organs were generated (Supp. Table 1). NHP tissues were processed in a Tissue-Tek VIP-6 vacuum infiltration processor (Sakura Finetek USA, Torrance, CA) followed by paraffin embedding with a Tissue-Tek model TEC (Sakura). Sections were cut on a Leica model 2245 microtome at 4 μm, stained with hematoxylin and eosin (H&E) and coverslipped. Slides were examined by an ACVP diplomate veterinary pathologist blinded to intervention. All images were captured with a Leica DM3000 microscope and DFC 500 digital camera using Leica Application Suite version 4.10.0 (Leica Microsystems, Buffalo Grove, IL).

### *In Situ* Hybridization

*In situ* hybridization (ISH) was performed using the RNAscope 2.5 HD RED kit (Advanced Cell Diagnostics, Newark, CA, USA) according to the manufacturer’s instructions. Briefly, EEEV ISH probe targeting nucleotides 8680-9901 of EEEV isolate V105-00210 was designed and synthesized by Advanced Cell Diagnostics (Cat# 455721). Tissue sections were deparaffinized with Xyless II (Valtech, Brackenridge, PA, USA), followed by a series of ethanol washes and peroxidase blocking, then heated in kit-provided antigen retrieval buffer, and digested by kit-provided proteinase. Sections were exposed to ISH target probe pairs and incubated at 40 °C in a hybridization oven for 2 h. After rinsing with wash buffer, ISH signal was amplified using kit-provided Pre-amplifier and Amplifier conjugated to alkaline phosphatase and incubated with Fast Red substrate solution for 10 mins at room temperature. Sections were then stained with hematoxylin, air-dried, and mounted. ISH images were collected using an Olympus BX53 upright microscope (Olympus Scientific Solutions Americas Corp., Waltham, MA, USA).

### Immunohistochemistry

Immunohistochemistry (IHC) was performed using the Dako Envision system (Dako Agilent Pathology Solutions, Carpinteria, CA, USA). After deparaffinization and peroxidase blocking, sections were covered with Rabbit anti-alphavirus polyclonal antibody (USAMRIID) at a dilution of 1:5000 and incubated at room temperature for 30 minutes. They were rinsed, and treated sequentially by an HRP-conjugated, secondary anti-rabbit polymer (Cat. #K4003, Dako Agilent Pathology Solutions). All slides were exposed to brown chromogenic substrate DAB (Cat. #K3468, Dako Agilent Pathology Solutions), counterstained with hematoxylin, dehydrated, cleared, and coverslipped. IHC images were collected using an Olympus BX53 upright microscope (Olympus Scientific Solutions Americas Corp., Waltham, MA, USA).

### Immunofluorescence Assay

Formalin-fixed paraffin embedded (FFPE) tissue sections were deparaffinized using xylene and a series of ethanol washes. After 0.1% Sudan black B (Sigma) treatment to eliminate the autofluorescence background, the sections were heated in Tris-EDTA buffer (10mM Tris Base, 1mM EDTA Solution, 0.05% Tween 20, pH 9.0) for 15 minutes to reverse formaldehyde crosslinks. After rinses with PBS (pH 7.4), the sections were blocked with PBS containing 5% normal goat serum overnight at 4°C. Then the sections were incubated with Rabbit anti-EEEV antibody (USAMRIID, 1:1000) and chicken anti-NeuN antibody (Abcam, 1:25), or chicken anti-GFAP (Abcam, 1:500, or mouse anti-CD68 (Agilent/Dako, 1:200) for 2 hours at room temperature. After rinses with PBS, the sections were incubated with secondary goat anti-chick Alex Fluor 488 (green, 1:500), goat anti-rabbit Alex Flour 488 (green), goat anti-rabbit Cy3 (red), and/ or goat anti-mouse Cy3 antibodies (red, 1:500) for 1 hour at room temperature. Sections were cover slipped using the Vectashield mounting medium with DAPI (Vector Laboratories). Images were captured on an LSM 880 Confocal Microscope (Zeiss, Oberkochen, Germany) and processed using open-source ImageJ software (National Institutes of Health, Bethesda, MD, USA).

### Transmission Electron Microscopy

Formalin-fixed thalamic tissue from each NHP was obtained and submerged in 2.5% glutaraldehyde and 2% paraformaldehyde in 0.1M sodium phosphate buffer for further fixation. Samples were fixed for at least 24 hours at 4°C and then rinsed with milliQ-EM grade water, rinsed again with 0.1M sodium cacodylate buffer before post-fixing with 1% osmium tetroxide in 0.1M sodium cacodylate for 60 minutes. After osmium fixation, the samples were rinsed with 0.1M sodium cacodylate buffer, followed by a water wash then subjected to uranyl acetate *en bloc*. Samples were washed with water then dehydrated through a graded ethanol series including 3 exchanges with 100% ethanol. Samples were further dehydrated with equal volumes of 100% ethanol and propylene oxide followed by two changes of propylene oxide. Samples were initially infiltrated with equal volumes of propylene oxide and resin (Embed-812; EMS, Hatfield, PA, USA) then incubated overnight in propylene oxide and resin. The next day, the samples were infiltrated with 100% resin embedded and oriented in 100% resin and then allowed to polymerize for 48 hours at 60 C. 1 micron thick sections were cut from one tissue block and a region of interest for thin sectioning was chosen. 80nm thin sections were cut and collected on 200 mesh copper grids. Two grids from each sample was further contrast stained with 2% uranyl acetate and Reynold’s lead citrate. Samples were then imaged on the Jeol 1011 TEM at various magnifications.

## Author Contributions

**Conceptualization:** Farooq Nasar, and Margaret L Pitt.

**Formal analysis:** Janice A Williams, Simon Y Long, Xiankun Zeng, John C

Trefry, Sharon Daye, Paul R Facemire, Patrick L Iversen, Sina Bavari, Margaret L Pitt, and Farooq Nasar.

**Funding acquisition:** Farooq Nasar and Margaret L Pitt.

**Investigation:** Janice A Williams, Simon Y Long, Xiankun Zeng, John C Trefry, Sharon Daye, Sina Bavari, Margaret L Pitt, and Farooq Nasar.

**Methodology:** Janice A Williams, Simon Y Long, Xiankun Zeng, Kathleen Kuehl, April M Babka, Neil M Davis, Jun Liu, and Sharon Daye.

**Project administration:** Farooq Nasar, and Margaret L. Pitt.

**Supervision:** Farooq Nasar, Sina Bavari, and Margaret L. Pitt.

**Writing – original draft**: Janice A Williams, Simon Y Long, and Farooq Nasar.

**Writing – review & editing**: Janice A Williams, Simon Y Long, Xiankun Zeng, Kathleen Kuehl, April M Babka, Neil M Davis, Jun Liu, John C Trefry, Sharon Daye, Paul R Facemire, Patrick L Iversen, Sina Bavari, Margaret L Pitt, and Farooq Nasar.

---

**Figure 22.** Proposed model of EEEV dissemination in the central nervous system (CNS) following an aerosol infection in cynomolgus macaques.

**Supp. Figure 1.** The absence of EEEV RNA in visceral organs of infected cynomolgus macaques. The tissues were collected at the time of euthanasia. The presence of viral RNA was visualized via *in situ* hybridization (ISH). ISH was performed on the tissues of all four NHPs. Bar = 200 um.

**Supp. Figure 2.** The absence of EEEV proteins in visceral organs of infected cynomolgus macaques. The tissues were collected at the time of euthanasia. The presence of viral proteins was visualized via immunohistochemistry (IHC). IHC was performed on the tissues of all four NHPs. Bar = 200 um.

**Supp. Figure 3.** The extracellular distribution of EEEV virions in the thalamus of infected cynomolgus macaques. Sections from NHPs were examined via transmission electron microscopy (TEM). Representative micrographs from each NHP are shown.

**Supp. Figure 4.** The localization of EEEV virions near synapses via transmission electron microscopy (TEM). Sections from the thalamus of each NHP were examined and representative micrographs from each NHP are shown. NHP #1 (A), NHP #2 (B), NHP #3 (C), and NHP #4 (D). Blue and red arrows show synapses and infectious virus particles, respectively. Bar = 600 nm.

**Supp. Figure 5.** Transmission electron microscopy (TEM) micrographs of viral replication centers within the brain of non-human primates. Top panels are representative electron micrographs of viral replication center (red asterisk) visible within the thalamus (A, E), amygdala (B, F), hippocampus (C, G) and hypothalamus (D, H) of a female non-human primate. The lower panels are also representative micrographs of the replication center in a male non-human primate. The number, size and intracellular localization of the replication center varies. A and F scale bar = 500 nm. B-E, G and H scale bar = 1 um.

**Supp. Table 1.** List of tissue sections from each organ.

## REFERENCES

1. Weston JH, Welsh MD, McLoughlin MF, Todd D. Salmon pancreas disease virus, an alphavirus infecting farmed Atlantic salmon, Salmo salar L. Virology. 1999;256(2):188–95. doi:10.1006/viro.1999.9654. PubMed PMID:10191183.

2. Villoing S, Bearzotti M, Chilmonczyk S, Castric J, Bremont M. Rainbow trout sleeping disease virus is an atypical alphavirus. J Virol. 2000;74(1):173–83. PubMed PMID:10590104; PubMed Central PMCID:PMCPMC111526.

3. Nasar F, Palacios G, Gorchakov RV, Guzman H, Da Rosa AP, Savji N, et al. Eilat virus, a unique alphavirus with host range restricted to insects by RNA replication. Proc Natl Acad Sci U S A. 2012;109(36):14622–7. doi:10.1073/pnas.1204787109. PubMed PMID:22908261; PubMed Central PMCID:PMCPMC3437828.

4. La Linn M, Gardner J, Warrilow D, Darnell GA, McMahon CR, Field I, et al. Arbovirus of marine mammals: a new alphavirus isolated from the elephant seal louse, Lepidophthirus macrorhini. J Virol. 2001;75(9):4103–9. doi:10.1128/JVI.75.9.4103-4109.2001. PubMed PMID:11287559; PubMed Central PMCID:PMCPMC114155.

5. Forrester NL, Palacios G, Tesh RB, Savji N, Guzman H, Sherman M, et al. Genome-scale phylogeny of the alphavirus genus suggests a marine origin. J Virol. 2012;86(5):2729–38. doi:10.1128/JVI.05591-11. PubMed PMID:22190718; PubMed Central PMCID:PMCPMC3302268.

6. Fields BN, Knipe DM, Howley PM. Fields virology. 6th ed. Philadelphia: Wolters Kluwer Health/Lippincott Williams & Wilkins; 2013.

7. Lindsey NP, Staples JE, Fischer M. Eastern Equine Encephalitis Virus in the United States, 2003-2016. Am J Trop Med Hyg. 2018;98(5):1472–7. Epub 2018/03/21. doi:10.4269/ajtmh.17-0927. PubMed PMID:29557336; PubMed Central PMCID:PMCPMC5953388.

8. Ko SY, Akahata W, Yang ES, Kong WP, Burke CW, Honnold SP, et al. A viruslike particle vaccine prevents equine encephalitis virus infection in nonhuman primates. Sci Transl Med. 2019;11(492). Epub 2019/05/17. doi:10.1126/scitranslmed.aav3113. PubMed PMID:31092692.

9. Reed DS, Lackemeyer MG, Garza NL, Norris S, Gamble S, Sullivan LJ, et al. Severe encephalitis in cynomolgus macaques exposed to aerosolized Eastern equine encephalitis virus. J Infect Dis. 2007;196(3):441–50. doi:10.1086/519391. PubMed PMID:17597459.

10. Roy CJ, Adams AP, Wang E, Leal G, Seymour RL, Sivasubramani SK, et al. A chimeric Sindbis-based vaccine protects cynomolgus macaques against a lethal aerosol challenge of eastern equine encephalitis virus. Vaccine. 2013;31(11):1464–70. doi:10.1016/j.vaccine.2013.01.014. PubMed PMID:23333212; PubMed Central PMCID:PMC3581708.

11. Porter AI, Erwin-Cohen RA, Twenhafel N, Chance T, Yee SB, Kern SJ, et al. Characterization and pathogenesis of aerosolized eastern equine encephalitis in the common marmoset (Callithrix jacchus). Virol J. 2017;14(1):25. Epub 2017/02/09. doi:10.1186/s12985-017-0687-7. PubMed PMID:28173871; PubMed Central PMCID:PMCPMC5297202.

12. Phelps AL, O’Brien LM, Eastaugh LS, Davies C, Lever MS, Ennis J, et al. Aerosol infection of Balb/c mice with eastern equine encephalitis virus; susceptibility and lethality. Virol J. 2019;16(1):2. Epub 2019/01/07. doi:10.1186/s12985-018-1103-7. PubMed PMID:30611287; PubMed Central PMCID:PMCPMC6321726.

13. Honnold SP, Mossel EC, Bakken RR, Lind CM, Cohen JW, Eccleston LT, et al. Eastern equine encephalitis virus in mice II: pathogenesis is dependent on route of exposure. Virol J. 2015;12:154. Epub 2015/10/02. doi:10.1186/s12985-015-0385-2. PubMed PMID:26423229; PubMed Central PMCID:PMCPMC4589026.

14. Roy CJ, Reed DS, Wilhelmsen CL, Hartings J, Norris S, Steele KE. Pathogenesis of aerosolized Eastern Equine Encephalitis virus infection in guinea pigs. Virol J. 2009;6:170. doi:10.1186/1743-422X-6-170. PubMed PMID:19852817; PubMed Central PMCID:PMC2770496.

15. Andrews C, Gerdin J, Patterson J, Buckles EL, Fitzgerald SD. Eastern equine encephalitis in puppies in Michigan and New York states. J Vet Diagn Invest. 2018;30(4):633–6. Epub 2018/05/03. doi:10.1177/1040638718774616. PubMed PMID:29717641; PubMed Central PMCID:PMCPMC6505920.

16. Bastian FO, Wende RD, Singer DB, Zeller RS. Eastern equine encephalomyelitis. Histopathologic and ultrastructural changes with isolation of the virus in a human case. Am J Clin Pathol. 1975;64(1):10–3. Epub 1975/07/01. doi:10.1093/ajcp/64.1.10. PubMed PMID:1171613.

17. Bauer RW, Gill MS, Poston RP, Kim DY. Naturally occurring eastern equine encephalitis in a Hampshire wether. J Vet Diagn Invest. 2005;17(3):281–5. Epub 2005/06/11. doi:10.1177/104063870501700314. PubMed PMID:15945389.

18. Belle EA, Grant LS, Thorburn MJ. An Outbreak of Eastern Equine Encephalomyelitis in Jamaica. Ii. Laboratory Diagnosis and Pathology of Eastern Equine Encephalomyelitis in Jamaica. Am J Trop Med Hyg. 1964;13:335–41. Epub 1964/03/01. doi:10.4269/ajtmh.1964.13.335. PubMed PMID:14125889.

19. Berlin D, Gilani AI, Grewal AK, Fowkes M. Eastern equine encephalitis. Pract Neurol. 2017;17(5):387–91. Epub 2017/07/30. doi:10.1136/practneurol-2017-001659. PubMed PMID:28754695.

20. Chenier S, Cote G, Vanderstock J, Macieira S, Laperle A, Helie P. An eastern equine encephalomyelitis (EEE) outbreak in Quebec in the fall of 2008. Can Vet J. 2010;51(9):1011–5. Epub 2010/12/02. PubMed PMID:21119870; PubMed Central PMCID:PMCPMC2920158.

21. Del Piero F, Wilkins PA, Dubovi EJ, Biolatti B, Cantile C. Clinical, pathologic, immunohistochemical, and virologic findings of eastern equine encephalomyelitis in two horses. Vet Pathol. 2001;38(4):451–6. Epub 2001/07/27. doi:10.1354/vp.38-4-451. PubMed PMID:11467481.

22. Deresiewicz RL, Thaler SJ, Hsu L, Zamani AA. Clinical and neuroradiographic manifestations of eastern equine encephalitis. N Engl J Med. 1997;336(26):1867–74. Epub 1997/06/26. doi:10.1056/NEJM199706263362604. PubMed PMID:9197215.

23. Elvinger F, Baldwin CA, Liggett AD, Tang KN, Dove CR. Protection of pigs by vaccination of pregnant sows against eastern equine encephalomyelitis virus. Vet Microbiol. 1996;51(3-4):229–39. Epub 1996/08/01. doi:10.1016/0378-1135(96)00037-5. PubMed PMID:8870186; PubMed Central PMCID:PMCPMC7117144.

24. Elvinger F, Liggett AD, Tang KN, Harrison LR, Cole JR, Jr., Baldwin CA, et al. Eastern equine encephalomyelitis virus infection in swine. J Am Vet Med Assoc. 1994;205(7):1014–6. Epub 1994/10/01. PubMed PMID:7852154.

25. Ethier M, Rogg J. Eastern equine encephalitis: MRI findings in two patients. Med Health R I. 2012;95(7):227–9. Epub 2012/08/30. PubMed PMID:22928238.

26. Farber S, Hill A, Connerly ML, Dingle JH. Encephalitis in infants and children - Caused by the virus of the eastern variety of equine encephalitis. Journal of the American Medical Association. 1940;114:1725–31. PubMed PMID:WOS:000201651800058.

27. Farrar MD, Miller DL, Baldwin CA, Stiver SL, Hall CL. Eastern equine encephalitis in dogs. J Vet Diagn Invest. 2005;17(6):614–7. Epub 2006/02/16. doi:10.1177/104063870501700619. PubMed PMID:16475527.

28. Feemster RF. Outbreak of Encephalitis in Man Due to the Eastern Virus of Equine Encephalomyelitis. Am J Public Health Nations Health. 1938;28(12):1403–10. Epub 1938/12/01. doi:10.2105/ajph.28.12.1403. PubMed PMID:18014957; PubMed Central PMCID:PMCPMC1527806.

29. Garen PD, Tsai TF, Powers JM. Human eastern equine encephalitis: immunohistochemistry and ultrastructure. Mod Pathol. 1999;12(6):646–52. Epub 1999/07/07. PubMed PMID:10392642.

30. Gregory CR, Latimer KS, Niagro FD, Campagnoli RP, Steffens WL, Ritchie BW. Detection of eastern equine encephalomyelitis virus RNA in formalin-fixed, paraffin-embedded tissues using DNA in situ hybridization. J Vet Diagn Invest. 1996;8(2):151–5. Epub 1996/04/01. doi:10.1177/104063879600800202. PubMed PMID:8744734.

31. Harvala H, Bremner J, Kealey S, Weller B, McLellan S, Lloyd G, et al. Case report: Eastern equine encephalitis virus imported to the UK. J Med Virol. 2009;81(2):305–8. Epub 2008/12/25. doi:10.1002/jmv.21379. PubMed PMID:19107960.

32. Hirsch MS, DeMaria A, Jr., Schaefer PW, Branda JA. Case records of the Massachusetts General Hospital. Case 22-2008. A 52-year-old woman with fever and confusion. N Engl J Med. 2008;359(3):294–303. Epub 2008/07/19. doi:10.1056/NEJMcpc0804149. PubMed PMID:18635435.

33. Hrabak T, Yerkey MW, Callerame K, Graham K. Eastern equine encephalitis presenting as psychosis. J Miss State Med Assoc. 2002;43(4):109–10. Epub 2002/05/07. PubMed PMID:11989192.

34. Jordan RA, Wagner JA, McCrumb FR. Eastern Equine Encephalitis: Report of a Case with Autopsy. Am J Trop Med Hyg. 1965;14:470–4. Epub 1965/05/01. doi:10.4269/ajtmh.1965.14.470. PubMed PMID:14292755.

35. Kim AS, Austin SK, Gardner CL, Zuiani A, Reed DS, Trobaugh DW, et al. Protective antibodies against Eastern equine encephalitis virus bind to epitopes in domains A and B of the E2 glycoprotein. Nat Microbiol. 2019;4(1):187–97. Epub 2018/11/21. doi:10.1038/s41564-018-0286-4. PubMed PMID:30455470; PubMed Central PMCID:PMCPMC6294662.

36. Kim JH, Booss J, Manuelidis EE, Duncan CC. Human eastern equine encephalitis. Electron microscopic study of a brain biopsy. Am J Clin Pathol. 1985;84(2):223–7. Epub 1985/08/01. doi:10.1093/ajcp/84.2.223. PubMed PMID:4025229.

37. Kiupel M, Fitzgerald SD, Pennick KE, Cooley TM, O’Brien DJ, Bolin SR, et al. Distribution of eastern equine encephalomyelitis viral protein and nucleic acid within central nervous tissue lesions in white-tailed deer (Odocoileus virginianus). Vet Pathol. 2013;50(6):1058–62. Epub 2013/05/21. doi:10.1177/0300985813488956. PubMed PMID:23686767.

38. Lury KM, Castillo M. Eastern equine encephalitis: CT and MRI findings in one case. Emerg Radiol. 2004;11(1):46–8. Epub 2004/08/17. doi:10.1007/s10140-004-0350-7. PubMed PMID:15309665.

39. McGee ED, Littleton CH, Mapp JB, Brown RJ. Eastern equine encephalomyelitis in an adult cow. Vet Pathol. 1992;29(4):361–3. Epub 1992/07/01. doi:10.1177/030098589202900414. PubMed PMID:1514224.

40. Morse RP, Bennish ML, Darras BT. Eastern equine encephalitis presenting with a focal brain lesion. Pediatr Neurol. 1992;8(6):473–5. Epub 1992/11/01. PubMed PMID:1476580.

41. Mukerji SS, Lam AD, Wilson MR. Eastern Equine Encephalitis Treated With Intravenous Immunoglobulins. Neurohospitalist. 2016;6(1):29–31. Epub 2016/01/08. doi:10.1177/1941874415578533. PubMed PMID:26740855; PubMed Central PMCID:PMCPMC4680893.

42. Nathanson N, Stolley PD, Boolukos PJ. Eastern equine encephalitis. Distribution of central nervous system lesions in man and Rhesus monkey. J Comp Pathol. 1969;79(1):109–15. Epub 1969/01/01. doi:10.1016/0021-9975(69)90034-6. PubMed PMID:4975613.

43. Nickerson JP, Kannabiran S, Burbank HN. MRI findings in eastern equine encephalitis: the “parenthesis” sign. Clin Imaging. 2016;40(2):222–3. Epub 2016/03/21. doi:10.1016/j.clinimag.2015.10.012. PubMed PMID:26995574.

44. Nolen-Walston R, Bedenice D, Rodriguez C, Rushton S, Bright A, Fecteau ME, et al. Eastern equine encephalitis in 9 South American camelids. J Vet Intern Med. 2007;21(4):846–52. Epub 2007/08/22. doi:10.1892/0891-6640(2007)21[846:eeeisa]2.0.co;2. PubMed PMID:17708408.

45. Patterson JS, Maes RK, Mullaney TP, Benson CL. Immunohistochemical diagnosis of eastern equine encephalomyelitis. J Vet Diagn Invest. 1996;8(2):156–60. Epub 1996/04/01. doi:10.1177/104063879600800203. PubMed PMID:8744735.

46. Pennick KE, McKnight CA, Patterson JS, Latimer KS, Maes RK, Wise AG, et al. Diagnostic sensitivity and specificity of in situ hybridization and immunohistochemistry for Eastern equine encephalitis virus and West Nile virus in formalin-fixed, paraffin-embedded brain tissue of horses. J Vet Diagn Invest. 2012;24(2):333–8. Epub 2012/03/02. doi:10.1177/1040638711435230. PubMed PMID:22379048.

47. Piliero PJ, Brody J, Zamani A, Deresiewicz RL. Eastern equine encephalitis presenting as focal neuroradiographic abnormalities: case report and review. Clin Infect Dis. 1994;18(6):985–8. Epub 1994/06/01. doi:10.1093/clinids/18.6.985. PubMed PMID:8086564.

48. Poonacha KB, Gregory CR, Vickers ML. Intestinal lesions in a horse associated with eastern equine encephalomyelitis virus infection. Vet Pathol. 1998;35(6):535–8. Epub 1998/11/21. doi:10.1177/030098589803500608. PubMed PMID:9823595.

49. Pouch SM, Katugaha SB, Shieh WJ, Annambhotla P, Walker WL, Basavaraju SV, et al. Transmission of Eastern Equine Encephalitis Virus from an Organ Donor to Three Transplant Recipients. Clin Infect Dis. 2018. Epub 2018/10/30. doi:10.1093/cid/ciy923. PubMed PMID:30371754; PubMed Central PMCID:PMCPMC6488434.

50. Przelomski MM, O’Rourke E, Grady GF, Berardi VP, Markley HG. Eastern equine encephalitis in Massachusetts: a report of 16 cases, 1970-1984. Neurology. 1988;38(5):736–9. Epub 1988/05/01. doi:10.1212/wnl.38.5.736. PubMed PMID:3362371.

51. Pursell AR, Mitchell FE, Seibold HR. Naturally occurring and experimentally induced eastern encephalomyelitis in calves. J Am Vet Med Assoc. 1976;169(10):1101–3. Epub 1976/11/15. PubMed PMID:977441.

52. Pursell AR, Peckham JC, Cole JR, Jr., Stewart WC, Mitchell FE. Naturally occurring and artificially induced eastern encephalomyelitis in pigs. J Am Vet Med Assoc. 1972;161(10):1143–7. Epub 1972/11/15. PubMed PMID:4652365.

53. Schmitt SM, Cooley TM, Fitzgerald SD, Bolin SR, Lim A, Schaefer SM, et al. An outbreak of Eastern equine encephalitis virus in free-ranging white-tailed deer in Michigan. J Wildl Dis. 2007;43(4):635–44. Epub 2007/11/07. doi:10.7589/0090-3558-43.4.635. PubMed PMID:17984258.

54. Shah KJ, Cherabuddi K. Case of eastern equine encephalitis presenting in winter. BMJ Case Rep. 2016;2016. Epub 2016/05/12. doi:10.1136/bcr-2016-215270. PubMed PMID:27165999; PubMed Central PMCID:PMCPMC4885365.

55. Silverman MA, Misasi J, Smole S, Feldman HA, Cohen AB, Santagata S, et al. Eastern equine encephalitis in children, Massachusetts and New Hampshire,USA, 1970-2010. Emerg Infect Dis. 2013;19(2):194–201; quiz 352. Epub 2013/01/25. doi:10.3201/eid1902.120039. PubMed PMID:23343480; PubMed Central PMCID:PMCPMC3559032.

56. Solomon IH, Ciarlini P, Santagata S, Ahmed AA, De Girolami U, Prasad S, et al. Fatal Eastern Equine Encephalitis in a Patient on Maintenance Rituximab: A Case Report. Open Forum Infect Dis. 2017;4(1):ofx021. Epub 2017/05/10. doi:10.1093/ofid/ofx021. PubMed PMID:28480291; PubMed Central PMCID:PMCPMC5414020.

57. Tate CM, Howerth EW, Stallknecht DE, Allison AB, Fischer JR, Mead DG. Eastern equine encephalitis in a free-ranging white-tailed deer (Odocoileus virginianus). J Wildl Dis. 2005;41(1):241–5. Epub 2005/04/14. doi:10.7589/0090-3558-41.1.241. PubMed PMID:15827230.

58. Vogel P, Kell WM, Fritz DL, Parker MD, Schoepp RJ. Early events in the pathogenesis of eastern equine encephalitis virus in mice. Am J Pathol. 2005;166(1):159–71. Epub 2005/01/06. doi:10.1016/S0002-9440(10)62241-9. PubMed PMID:15632009; PubMed Central PMCID:PMCPMC1602312.

59. Adams AP, Aronson JF, Tardif SD, Patterson JL, Brasky KM, Geiger R, et al. Common marmosets (Callithrix jacchus) as a nonhuman primate model to assess the virulence of eastern equine encephalitis virus strains. J Virol. 2008;82(18):9035–42. Epub 2008/07/11. doi:10.1128/JVI.00674-08. PubMed PMID:18614636; PubMed Central PMCID:PMCPMC2546911.

60. Hurst EW. Infection of therhesus monkey (Macaca mulatta) and the guinea-pig with the virus of equine encephalomyelitis. The Journal of Pathology and Bacteriology. 1936;42(1):271–302. doi:10.1002/path.1700420128.

61. Jungherr EL, Helmboldt CF, Satriano SF, Luginbuhl RE. Investigation of eastern equine encephalomyelitis. III. Pathology in pheasants and incidental observations in feral birds. Am J Hyg. 1958;67(1):10–20. Epub 1958/01/01. PubMed PMID:13508649.

62. King LS. Studies on Eastern Equine Encephalomyelitis: Ii. Pathogenesis of the Disease in the Guinea Pig. J Exp Med. 1939;69(5):675–90. Epub 1939/04/30. doi:10.1084/jem.69.5.675. PubMed PMID:19870870; PubMed Central PMCID:PMCPMC2133754.

63. King LS. Studies on Eastern Equine Encephalomyelitis: V. Histopathology in the Mouse. J Exp Med. 1940;71(1):107–12. Epub 1940/01/01. doi:10.1084/jem.71.1.107. PubMed PMID:19870938; PubMed Central PMCID:PMCPMC2134999.

64. Ognianov D, Fernandez A. [Studies on the pathogenesis of eastern equine encephalomyelitis (EEE) in the mouse after injections of virus]. Zentralbl Veterinarmed B. 1972;19(2):89–93. Epub 1972/02/01. PubMed PMID:5050029.

65. Paessler S, Aguilar P, Anishchenko M, Wang HQ, Aronson J, Campbell G, et al. The hamster as an animal model for eastern equine encephalitis--and its use in studies of virus entrance into the brain. J Infect Dis. 2004;189(11):2072–6. Epub 2004/05/15. doi:10.1086/383246. PubMed PMID:15143475.

66. Wendell LC, Potter NS, Roth JL, Salloway SP, Thompson BB. Successful management of severe neuroinvasive eastern equine encephalitis. Neurocrit Care. 2013;19(1):111–5. Epub 2013/06/05. doi:10.1007/s12028-013-9822-5. PubMed PMID:23733173.

67. Murphy FA, Whitfield SG. Eastern equine encephalitis virus infection: electron microscopic studies of mouse central nervous system. Exp Mol Pathol. 1970;13(2):131–46. Epub 1970/10/01. doi:10.1016/0014-4800(70)90001-8. PubMed PMID:5470808.

68. Zhang R, Hryc CF, Cong Y, Liu X, Jakana J, Gorchakov R, et al. 4.4 A cryo-EM structure of an enveloped alphavirus Venezuelan equine encephalitis virus. EMBO J. 2011;30(18):3854–63. Epub 2011/08/11. doi:10.1038/emboj.2011.261. PubMed PMID:21829169; PubMed Central PMCID:PMCPMC3173789.

69. Sherman MB, Weaver SC. Structure of the recombinant alphavirus Western equine encephalitis virus revealed by cryoelectron microscopy. J Virol. 2010;84(19):9775–82. Epub 2010/07/16. doi:10.1128/JVI.00876-10. PubMed PMID:20631130; PubMed Central PMCID:PMCPMC2937749.

70. Chen L, Wang M, Zhu D, Sun Z, Ma J, Wang J, et al. Implication for alphavirus host-cell entry and assembly indicated by a 3.5A resolution cryo-EM structure. Nat Commun. 2018;9(1):5326. Epub 2018/12/16. doi:10.1038/s41467-018-07704-x. PubMed PMID:30552337; PubMed Central PMCID:PMCPMC6294011.

71. Kostyuchenko VA, Jakana J, Liu X, Haddow AD, Aung M, Weaver SC, et al. The structure of barmah forest virus as revealed by cryo-electron microscopy at a 6-angstrom resolution has detailed transmembrane protein architecture and interactions. J Virol. 2011;85(18):9327–33. Epub 2011/07/15. doi:10.1128/JVI.05015-11. PubMed PMID:21752915; PubMed Central PMCID:PMCPMC3165765.

72. Boonstra H, Oosterhuis JW, Oosterhuis AM, Fleuren GJ. Cervical tissue shrinkage by formaldehyde fixation, paraffin wax embedding, section cutting and mounting. Virchows Arch A Pathol Anat Histopathol. 1983;402(2):195–201. Epub 1983/01/01. doi:10.1007/BF00695061. PubMed PMID:6420986.

73. Luft JH. Embedding Media — Old and New. Advanced Techniques in Biological Electron Microscopy 1973. p. 1–34.

74. Crang RFE, Klomparens KL. Artifacts in Biological Electron Microscopy: Springer; 1988.

75. Maday S, Twelvetrees AE, Moughamian AJ, Holzbaur EL. Axonal transport: cargo-specific mechanisms of motility and regulation. Neuron. 2014;84(2):292–309. Epub 2014/11/07. doi:10.1016/j.neuron.2014.10.019. PubMed PMID:25374356; PubMed Central PMCID:PMCPMC4269290.

76. Monath TP, Cropp CB, Harrison AK. Mode of entry of a neurotropic arbovirus into the central nervous system. Reinvestigation of an old controversy. Lab Invest. 1983;48(4):399–410. Epub 1983/04/01. PubMed PMID:6300550.

77. Bauer A, Nolden T, Schroter J, Romer-Oberdorfer A, Gluska S, Perlson E, et al. Anterograde glycoprotein-dependent transport of newly generated rabies virus in dorsal root ganglion neurons. J Virol. 2014;88(24):14172–83. Epub 2014/10/03. doi:10.1128/JVI.02254-14. PubMed PMID:25275124; PubMed Central PMCID:PMCPMC4249153.

78. Samuel MA, Wang H, Siddharthan V, Morrey JD, Diamond MS. Axonal transport mediates West Nile virus entry into the central nervous system and induces acute flaccid paralysis. Proc Natl Acad Sci U S A. 2007;104(43):17140–5. Epub 2007/10/18. doi:10.1073/pnas.0705837104. PubMed PMID:17939996; PubMed Central PMCID:PMCPMC2040476.

79. Qin C, Li J, Tang K. The Paraventricular Nucleus of the Hypothalamus: Development, Function, and Human Diseases. Endocrinology. 2018;159(9):3458–72. Epub 2018/07/28. doi:10.1210/en.2018-00453. PubMed PMID:30052854.

80. Lozic M, Sarenac O, Murphy D, Japundzic-Zigon N. Vasopressin, Central Autonomic Control and Blood Pressure Regulation. Curr Hypertens Rep. 2018;20(2):11. Epub 2018/02/27. doi:10.1007/s11906-018-0811-0. PubMed PMID:29480411.

81. Herzog ED, Hermanstyne T, Smyllie NJ, Hastings MH. Regulating the Suprachiasmatic Nucleus (SCN) Circadian Clockwork: Interplay between Cell-Autonomous and Circuit-Level Mechanisms. Cold Spring Harb Perspect Biol. 2017;9(1). Epub 2017/01/05. doi:10.1101/cshperspect.a027706. PubMed PMID:28049647; PubMed Central PMCID:PMCPMC5204321.

82. Szymusiak R, McGinty D. Hypothalamic regulation of sleep and arousal. Ann N Y Acad Sci. 2008;1129:275–86. Epub 2008/07/02. doi:10.1196/annals.1417.027. PubMed PMID:18591488.

83. Fukushi I, Yokota S, Okada Y. The role of the hypothalamus in modulation of respiration. Respir Physiol Neurobiol. 2019;265:172–9. Epub 2018/07/17. doi:10.1016/j.resp.2018.07.003. PubMed PMID:30009993.

84. Rahmouni K. Cardiovascular Regulation by the Arcuate Nucleus of the Hypothalamus: Neurocircuitry and Signaling Systems. Hypertension. 2016;67(6):1064–71. Epub 2016/04/06. doi:10.1161/HYPERTENSIONAHA.115.06425. PubMed PMID:27045026; PubMed Central PMCID:PMCPMC4865428.

85. Zimmerman CA, Leib DE, Knight ZA. Neural circuits underlying thirst and fluid homeostasis. Nat Rev Neurosci. 2017;18(8):459–69. Epub 2017/06/24. doi:10.1038/nrn.2017.71. PubMed PMID:28638120; PubMed Central PMCID:PMCPMC5955721.

86. Tan CL, Knight ZA. Regulation of Body Temperature by the Nervous System. Neuron. 2018;98(1):31–48. Epub 2018/04/06. doi:10.1016/j.neuron.2018.02.022. PubMed PMID:29621489; PubMed Central PMCID:PMCPMC6034117.

87. Fu ZF, Weihe E, Zheng YM, Schafer MK, Sheng H, Corisdeo S, et al. Differential effects of rabies and borna disease viruses on immediate-early- and late-response gene expression in brain tissues. J Virol. 1993;67(11):6674–81. Epub 1993/11/01. PubMed PMID:8411369; PubMed Central PMCID:PMCPMC238106.

88. Tsiang H. Neuronal function impairment in rabies-infected rat brain. J Gen Virol. 1982;61 (Pt 2):277–81. Epub 1982/08/01. doi:10.1099/0022-1317-61-2-277. PubMed PMID:7119753.

89. Bouzamondo E, Ladogana A, Tsiang H. Alteration of potassium-evoked 5-HT release from virus-infected rat cortical synaptosomes. Neuroreport. 1993;4(5):555–8. Epub 1993/05/01. doi:10.1097/00001756-199305000-00023. PubMed PMID:8513137.

90. Ceccaldi PE, Fillion MP, Ermine A, Tsiang H, Fillion G. Rabies virus selectively alters 5-HT1 receptor subtypes in rat brain. Eur J Pharmacol. 1993;245(2):129–38. Epub 1993/04/15. doi:10.1016/0922-4106(93)90120-x. PubMed PMID:8491253.

91. Iwata M, Unno T, Minamoto N, Ohashi H, Komori S. Rabies virus infection prevents the modulation by alpha(2)-adrenoceptors, but not muscarinic receptors, of Ca(2+) channels in NG108-15 cells. Eur J Pharmacol. 2000;404(1-2):79–88. Epub 2000/09/12. doi:10.1016/s0014-2999(00)00621-x. PubMed PMID:10980265.

92. Iwata M, Komori S, Unno T, Minamoto N, Ohashi H. Modification of membrane currents in mouse neuroblastoma cells following infection with rabies virus. Br J Pharmacol. 1999;126(8):1691–8. Epub 1999/06/18. doi:10.1038/sj.bjp.0702473. PubMed PMID:10372810; PubMed Central PMCID:PMCPMC1565954.

93. Scott CA, Rossiter JP, Andrew RD, Jackson AC. Structural abnormalities in neurons are sufficient to explain the clinical disease and fatal outcome of experimental rabies in yellow fluorescent protein-expressing transgenic mice. J Virol. 2008;82(1):513–21. Epub 2007/10/19. doi:10.1128/JVI.01677-07. PubMed PMID:17942540; PubMed Central PMCID:PMCPMC2224401.

94. Sundaramoorthy V, Green D, Locke K, O’Brien CM, Dearnley M, Bingham J. Novel role of SARM1 mediated axonal degeneration in the pathogenesis of rabies. PLoS Pathog. 2020;16(2):e1008343. Epub 2020/02/19. doi:10.1371/journal.ppat.1008343. PubMed PMID:32069324; PubMed Central PMCID:PMCPMC7048299.

95. National Research Council (U.S.). Committee for the Update of the Guide for the Care and Use of Laboratory Animals., Institute for Laboratory Animal Research (U.S.), National Academies Press (U.S.). Guide for the care and use of laboratory animals. 8th ed. Washington, D.C.: National Academies Press; 2011. xxv, 220 p. p.

