## Supplementary material for "Eastern Equine Encephalitis Virus Rapidly Infects and Disseminates in the Brain and Spinal Cord of Infected Cynomolgus Macaques Following Aerosol Challenge": Supp. Figs. 1-5, Table 1

### Supp. Figure 1

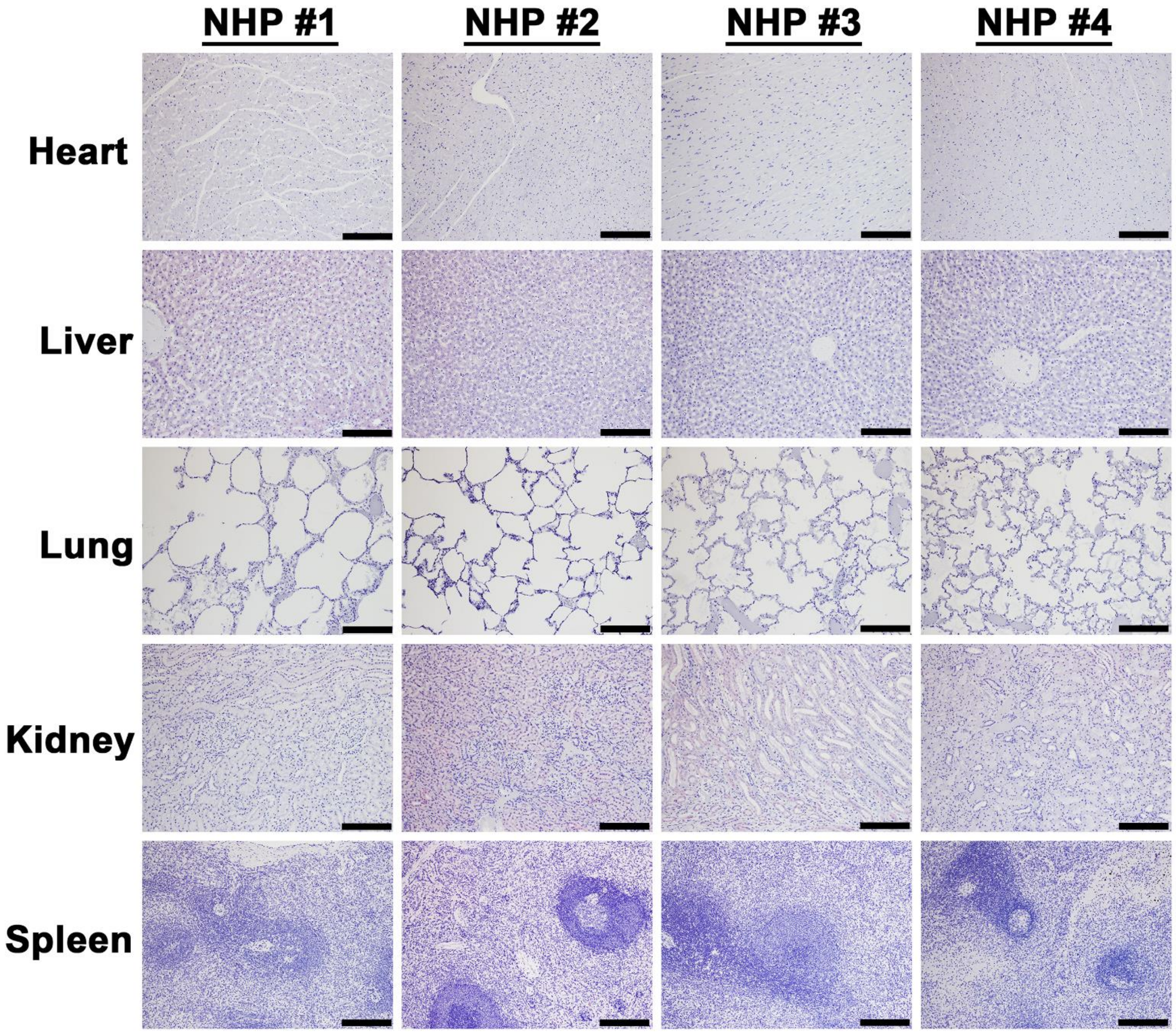

### Supp. Figure 2

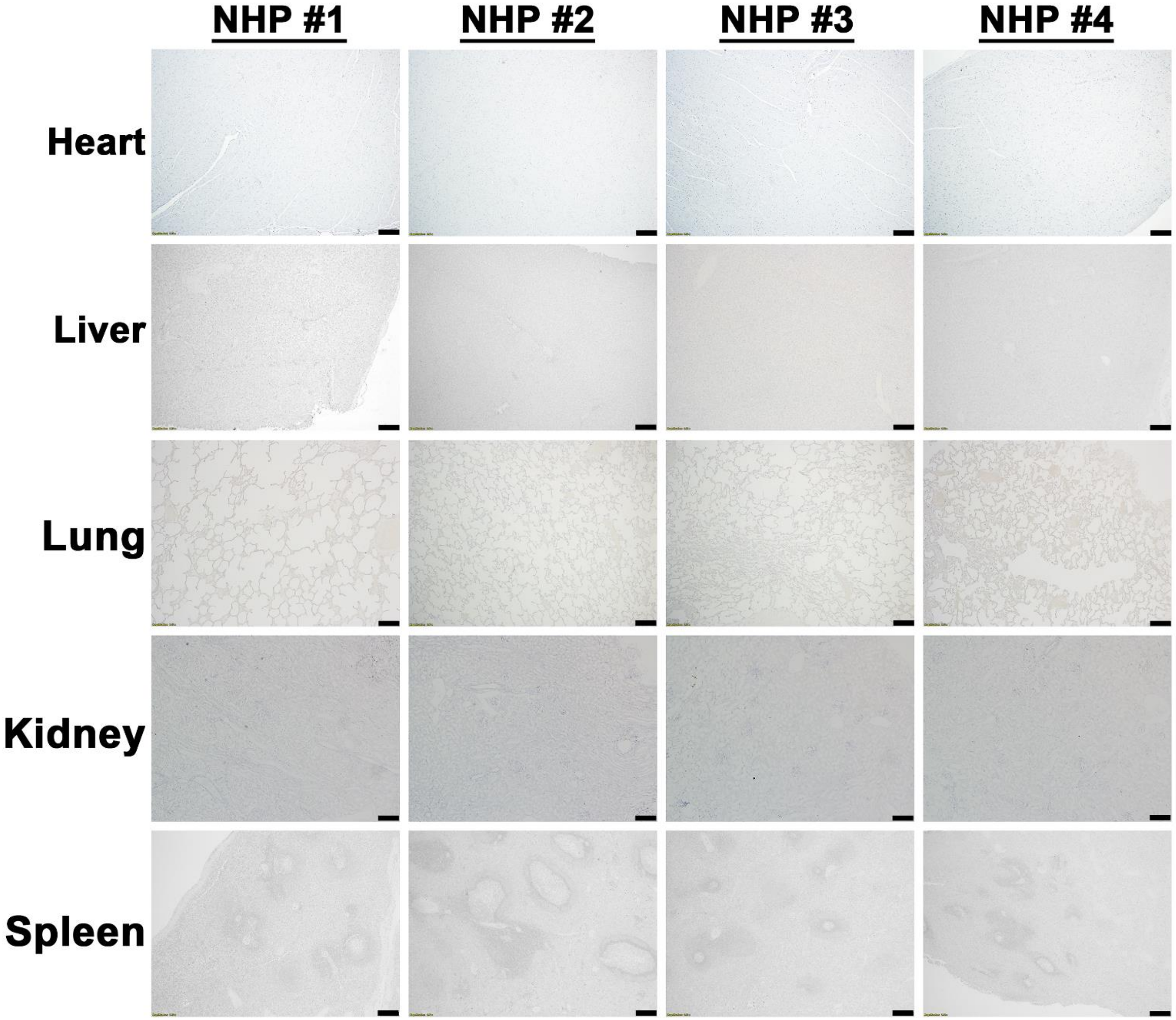

### Supp. Figure 3

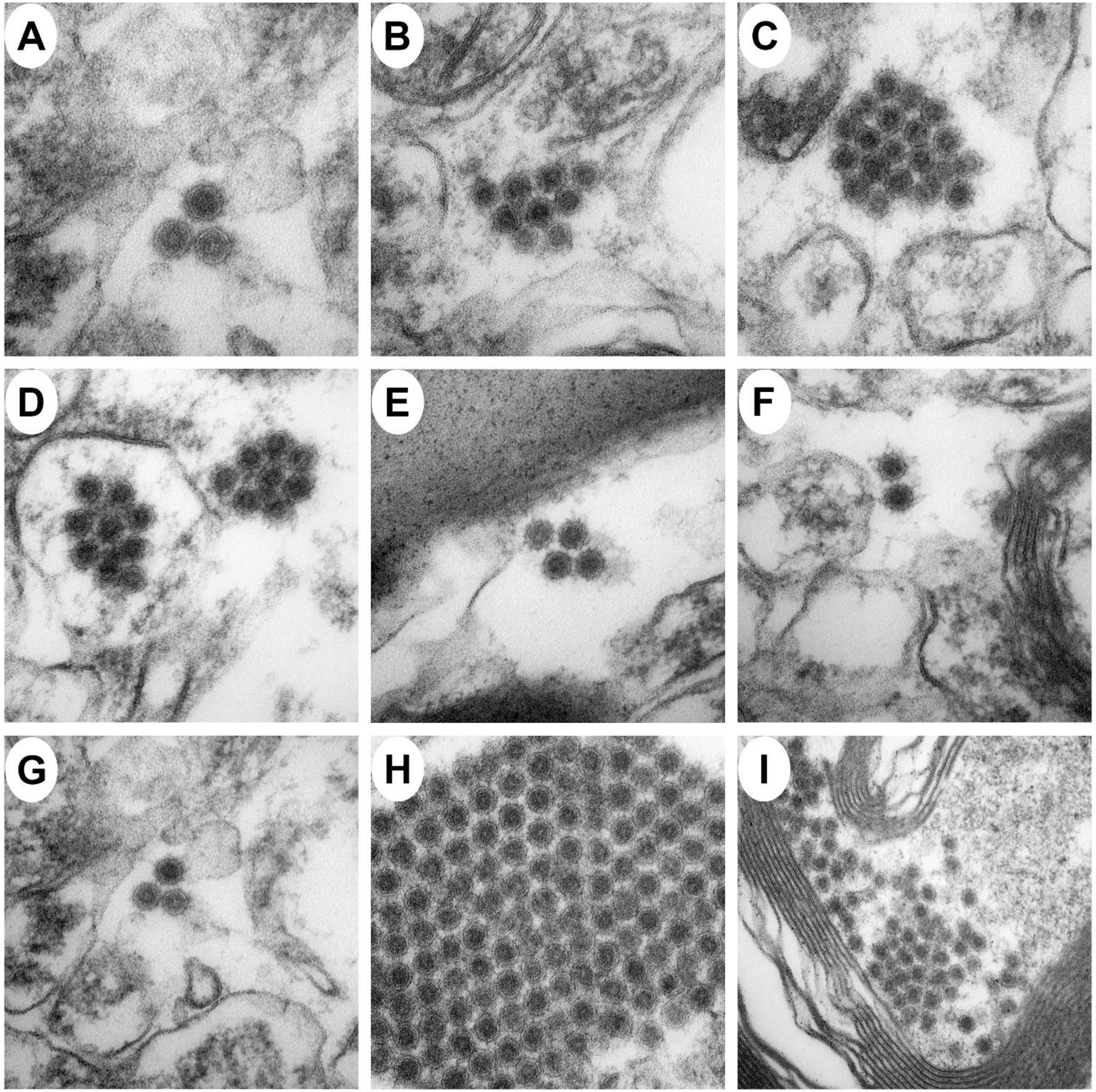

### Supp. Figure 4

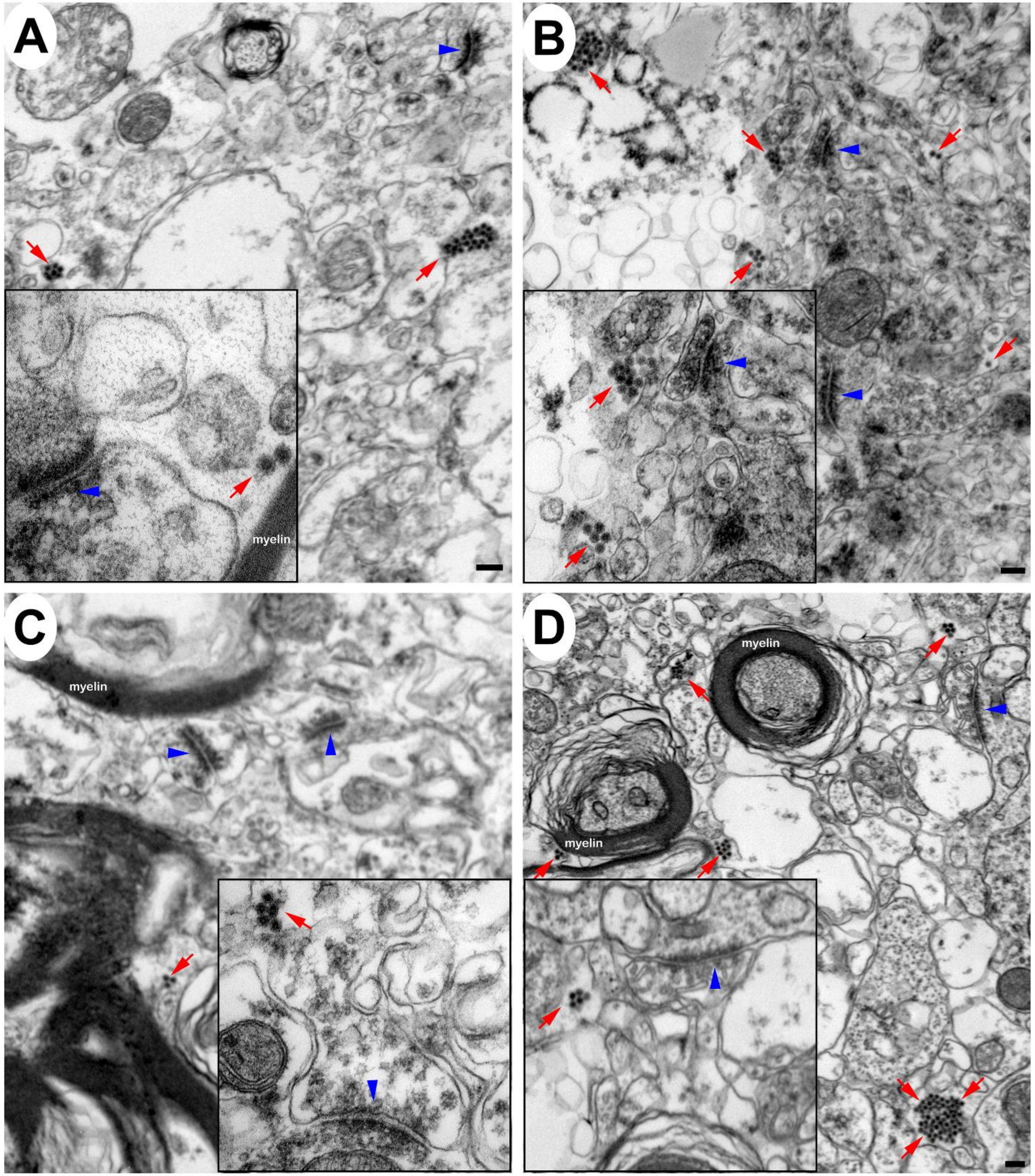

### Supp. Figure 5

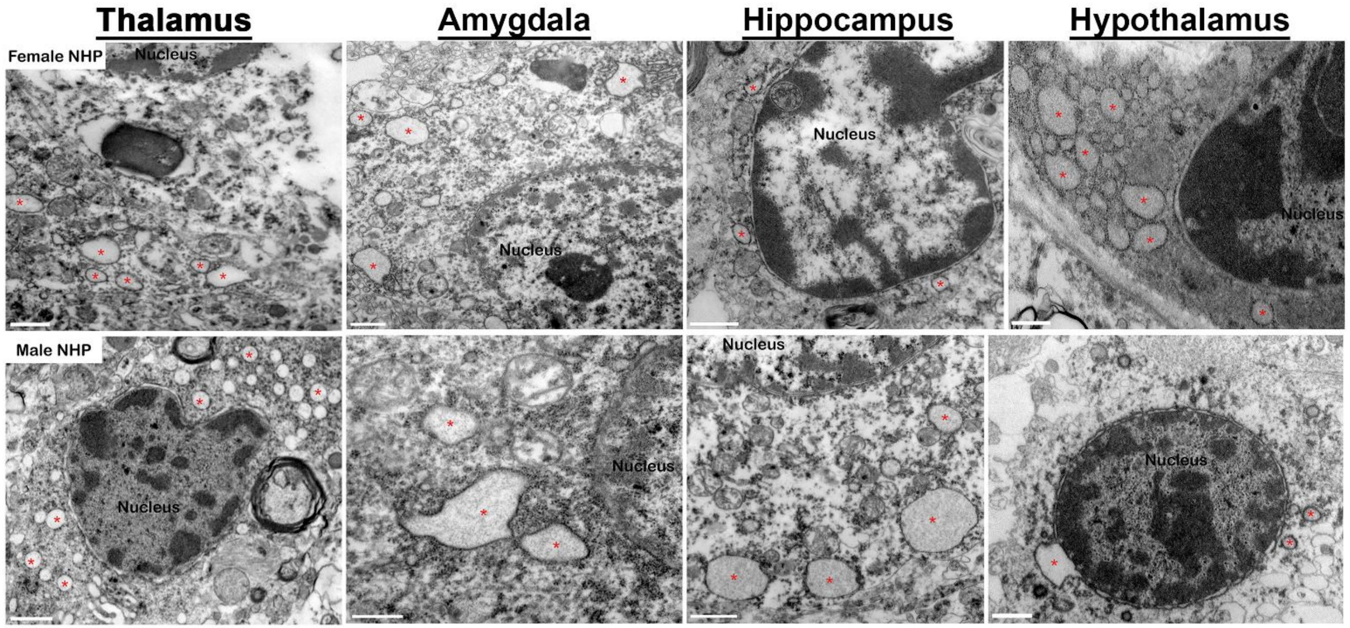

### Supp. Table 1

| Tissue | Tissue Sections |
|---|---|
| Lung | Right cranial lobe |
| | Left cranial lobe |
| | Right caudal lobe |
| | Left caudal lobe |
| Kidney | Right (transverse section) |
| | Left (longitudinal section) |
| Heart | Right ventricle |
| | Left ventricle |
| | Interventricular septum |
| Liver | Liver with gallbladder |
| | Right Lobe |
| | Left Lobe |
| Spleen | One cross section |
| Brain | Frontal Cortex |
| | Corpus Striatum |
| | Hypothalamus |
| | Thalamus |
| | Mesensephalon |
| | Medulla Oblongata |
| | Cerebellum |
| | Amygdala |
| | Hippocampus |
